# Rapid evolution of piRNA clusters in the *Drosophila melanogaster* ovary

**DOI:** 10.1101/2023.05.08.539910

**Authors:** Satyam Srivastav, Cédric Feschotte, Andrew G. Clark

## Abstract

Animal genomes are parasitized by a horde of transposable elements (TEs) whose mutagenic activity can have catastrophic consequences. The piRNA pathway is a conserved mechanism to repress TE activity in the germline via a specialized class of small RNAs associated with effector Piwi proteins called piwi-associated RNAs (piRNAs). piRNAs are produced from discrete genomic regions called piRNA clusters (piCs). While piCs are generally enriched for TE sequences and the molecular processes by which they are transcribed and regulated are relatively well understood in *Drosophila melanogaster*, much less is known about the origin and evolution of piCs in this or any other species. To investigate piC evolution, we use a population genomics approach to compare piC activity and sequence composition across 8 geographically distant strains of *D. melanogaster* with high quality long-read genome assemblies. We perform extensive annotations of ovary piCs and TE content in each strain and test predictions of two proposed models of piC evolution. The *‘de novo’* model posits that individual TE insertions can spontaneously attain the status of a small piC to generate piRNAs silencing the entire TE family. The ‘trap’ model envisions large and evolutionary stable genomic clusters where TEs tend to accumulate and serves as a long-term “memory” of ancient TE invasions and produce a great variety of piRNAs protecting against related TEs entering the genome. It remains unclear which model best describes the evolution of piCs. Our analysis uncovers extensive variation in piC activity across strains and signatures of rapid birth and death of piCs in natural populations. Most TE families inferred to be recently or currently active show an enrichment of strain-specific insertions into large piCs, consistent with the trap model. By contrast, only a small subset of active LTR retrotransposon families is enriched for the formation of strain-specific piCs, suggesting that these families have an inherent proclivity to form *de novo* piCs. Thus, our findings support aspects of both *‘de novo’* and ‘trap’ models of piC evolution. We propose that these two models represent two extreme stages along an evolutionary continuum, which begins with the emergence of piCs *de novo* from a few specific LTR retrotransposon insertions that subsequently expand by accretion of other TE insertions during evolution to form larger ‘trap’ clusters. Our study shows that piCs are evolutionarily labile and that TEs themselves are the major force driving the formation and evolution of piCs.

## Introduction

Organisms have evolved many mechanisms to minimize the genomic instability caused by transposable element (TE) activity (Bingham et al. 1982; Hedges and Deininger 2007; Huang et al. 2012; Montgomery et al. 1991). In animals, the Piwi-associated RNA (piRNA) pathway is a conserved small RNA-based mechanism regulating TE activity in the germline (Brennecke et al. 2007; Houwing et al. 2007; Lau et al. 2006; Grimson et al. 2008). piRNAs are 23-35 nucleotide RNAs produced from discrete loci called piRNA clusters (piCs) that guide Piwi effector proteins to silence TEs (Saito and Siomi 2010; Ozata et al. 2019). The piRNA pathway presents features of an adaptive defense system against TE invasion (Brennecke et al. 2008; Kofler et al. 2018; Khurana et al. 2011; Yu et al. 2019) but little is known about the processes and principles driving its evolution. The genes encoding the effector proteins and processing factors involved in piRNA-mediated silencing display signatures of adaptive evolution (positive selection) in several species’ lineages (Yi et al. 2014; Simkin et al. 2013; Vermaak et al. 2005; Palmer et al. 2018), which may indicate adaptation to rapidly changing TE sequences and new invasions (Cosby et al. 2019). While attempts have been made to explain the rapid evolution of piRNA pathway genes, (Wang et al. 2020; Parhad et al. 2020; Brand and Levine 2021) little is known about how piRNA-producing loci originate and evolve in flies, or any other species.

piRNAs are produced from single-stranded long non-coding RNA precursors that are transcribed from dispersed loci called piRNA clusters (Li et al. 2013; Mohn et al. 2014; Brennecke et al. 2007). piCs make up 0.1-3% of the genome of flies, mosquitoes, and mice and are enriched for TEs and other repeats such as DNA satellites but sometimes host gene sequences as well (Chirn et al. 2015; Brennecke et al. 2007; Ma et al. 2021; Roovers et al. 2015; Chen et al. 2021). The best characterized function of piRNAs is to repress TEs. Since TE activity and composition vary significantly between and within species, TEs themselves must be important drivers of piC evolution, but this has not been thoroughly tested. TEs exhibit high diversity in their mechanisms of transposition and genomic distribution (Sultana et al. 2017; Charlesworth et al. 1994; Bartolomé et al. 2002; Wells and Feschotte 2020). In addition, differences in the spatial and temporal activity of TE families exist in animal germlines (Laski et al. 1986; Calvi and Gelbart 1994; Bogu et al. 2019; Yoth et al. 2022; Chang et al. 2022). Hence, it is likely that piCs evolve through diverse mechanisms to repress newly introduced TEs, which is predicted to create an arms race between TEs and piCs (Cosby et al. 2019; Luo et al. 2020; Parhad and Theurkauf 2019; Said et al. 2022). Indeed, some piCs in flies are specialized to repress specific subsets of TE families, such as *flamenco*, which is almost entirely composed of and dedicated to silencing *Ty3/mdg4* (formerly known as *gypsy*) retroviral-like elements (Zanni et al. 2013). Although detailed mechanistic features of piC regulation and function have been uncovered in the last decade (Ozata et al. 2019; Czech et al. 2018), there is very limited understanding of piC evolution. Thus far, studies of piC evolution have been restricted to either a few large piCs or to small conserved genic piCs (Gebert et al. 2021; Chirn et al. 2015; Zhang et al. 2020; Ellison and Cao 2020; Mohamed et al. 2020; Wierzbicki et al. 2023).

The organization of piCs is best characterized in *Drosophila melanogaster*. The genome-wide piC landscape in *D. melanogaster* ovary comprises of tens of large (>10 kb) loci and hundreds of smaller (<10 kb) loci (Brennecke et al. 2007; Chen et al. 2021). It is also known that most large clusters (>10 kb) reside in pericentromeric and sub-telomeric regions. Larger pericentromeric clusters are composed of tens to hundreds of diverse TE insertions, while the small clusters (<10 kb) often contain recent TE insertions (Shpiz et al. 2014; Miller et al. 2023; Robine et al. 2009). The architecture and composition of some large clusters suggests a ‘trap’ model for the evolution of piCs, wherein TE insertions of active families within clusters is selectively favored because of their presumed transposition repressive effects (Zanni et al. 2013; Bergman et al. 2006; Moon et al. 2018; Zhang et al. 2020). Over time, this process is predicted to lead to the accumulation of archival remnants of past TE invasions in piCs. Thus, these large piCs are thought to produce a bank of diverse piRNAs related to previously encountered TEs as means to silence newly introduced TEs. On the other hand, the finding that small piCs can originate from recent TE insertions suggested a *‘de novo’* model. Here, individual TE insertions are converted into piCs through an epigenetic licensing process guided by maternally deposited piRNAs (le Thomas et al. 2014; Brennecke et al. 2008; Olovnikov et al. 2013). The ‘*de novo*’ model abrogates the need for active TEs to land into existing ‘trap’ clusters to come under the control of the piRNA pathway (Shpiz et al. 2014; Gebert et al. 2021).

The ‘trap’ and ‘*de novo*’ models of piC evolution are not mutually exclusive, but each makes contrasting predictions on how the structure, composition and activity of piCs are expected to evolve in a population. In the ‘trap’ model, piCs are expected to be (1) fewer in number, larger (10-100 kb), and mainly peri-centromeric and sub-telomeric; (2) piCs are expected to be syntenically stable archive of sequences of active and inactive TE families, and (3) piCs should gradually undergo TE sequence turnover via internal sequence rearrangements such as insertions and deletions (INDELs). The latter process is likely to be a fundamental characteristic of ‘traps’, wherein insertions of young and active TE families would replace older insertions of inactive TE families. That is because there is a limit to the genomic space afforded to clusters due to the possibility of spreading of their silent chromatin over host genes (Blumenstiel et al. 2016; Huang et al. 2022; Lee and Karpen 2017). This would also ensure that piC sequences are representative of the ever-changing genomic TE content while maintaining a constant genomic space over time (Kofler 2020; Said et al. 2022). Initial discoveries of piC architecture and composition in *D. melanogaster* were consistent with the ‘trap’ model (Assis and Kondrashov 2009; Brennecke et al. 2007; Malone et al. 2009). Notably, the *flamenco* cluster on the X chromosome is proposed to act as a ‘trap’ specialized for the capture of *Ty3*/*mdg4* retrotransposons (Genzor et al. 2021; Zanni et al. 2013). Furthermore, evolution of piRNA-mediated repression of recently invading TEs, such as the *P*-element, documented cases where silencing of *P*-elements is established by individual insertions within large known piCs – *42AB, #40, #3* and *X-TAS* both in laboratory and natural conditions (Khurana et al. 2011; Zhang et al. 2020; Moon et al. 2018; Ronsseray et al. 1991; Srivastav et al. 2019). While these studies suggest that horizontally acquired TEs come under the control of piRNA produced by insertions from the same family inserted into existing piCs, it remains unclear whether the ‘trap’ concept is broadly generalizable.

The ‘*de novo*’ model of piC evolution posits that TE silencing is driven by smaller clusters comprised of individual TE insertions of recent origin. If the ‘*de novo’* model is the primary mode of piC evolution, it should be expected that piCs are 1) abundant but smaller in size (1-10 kb), broadly distributed genome-wide, 2) highly polymorphic in population and typically strain-specific, and comprised mainly of insertions from active TE families, and 3) exhibit wholescale insertions and deletions, leading to loss or gain of entire clusters. These properties would ensure that piRNA production is biased toward active TE families. Several independent studies have provided observations that *de novo* formation of canonical piCs can emerge in the laboratory from artificial transgenes (Muerdter et al. 2012; de Vanssay et al. 2012) and in nature from recent TE insertions (Shpiz et al. 2014; Ryazansky et al. 2017). In summary, while both trap and *de novo* models of piC evolution have received empirical support, their relative contribution to piC evolution in *D. melanogaster* is unclear, and the mechanisms underlying the evolution of piCs remain broadly uncharacterized. In the present study, we found that there is extensive variation in sequence and activity of piCs across 8 *D. melanogaster* strains. These results lead us to develop a unified model of piC evolution that integrate components of *‘de novo’* and ‘trap’ models supported by our findings.

## Results

### Extensive variation in the genomic landscape of piCs

To quantify piC variation in *D. melanogaster* we generate a comprehensive annotation of active piCs in eight highly inbred strains. Seven of these strains are derived from natural populations of distinct worldwide origins and have publicly available long-read genome assemblies (Chakraborty et al. 2019). For each of these seven strains, we constructed and sequenced libraries of small RNAs isolated from ovaries of two biological replicates sampled 6 months apart (**Supplementary Fig. S1A, Table S1)** (see Methods). In addition, we analyzed two ovarian small RNA libraries for the reference iso-1 strain, generated as part of two independent studies (Shipz et al, 2014 and Asif-Laidin et al, 2017). piCs are defined by high expression and density of 23-29 nt long and 1U-biased small RNAs inferred to represent piRNAs (Brennecke et al. 2007; Mohn et al. 2014). Each small RNA library is analyzed separately using the pipeline outlined briefly here and described in more detail in Methods, which lists three methods to define piCs **(Fig. 1A, Supplementary Fig. 2B)**. The *restrictive* and *proTRAC* methods serve the purpose of discovering moderately to highly expressed piCs using uniquely and multi-mapping piRNAs respectively. The *permissive* method is carried out mainly to validate low to moderately expressed piCs detected by *proTRAC* using multi-mapping piRNAs.

**Figure 1.**
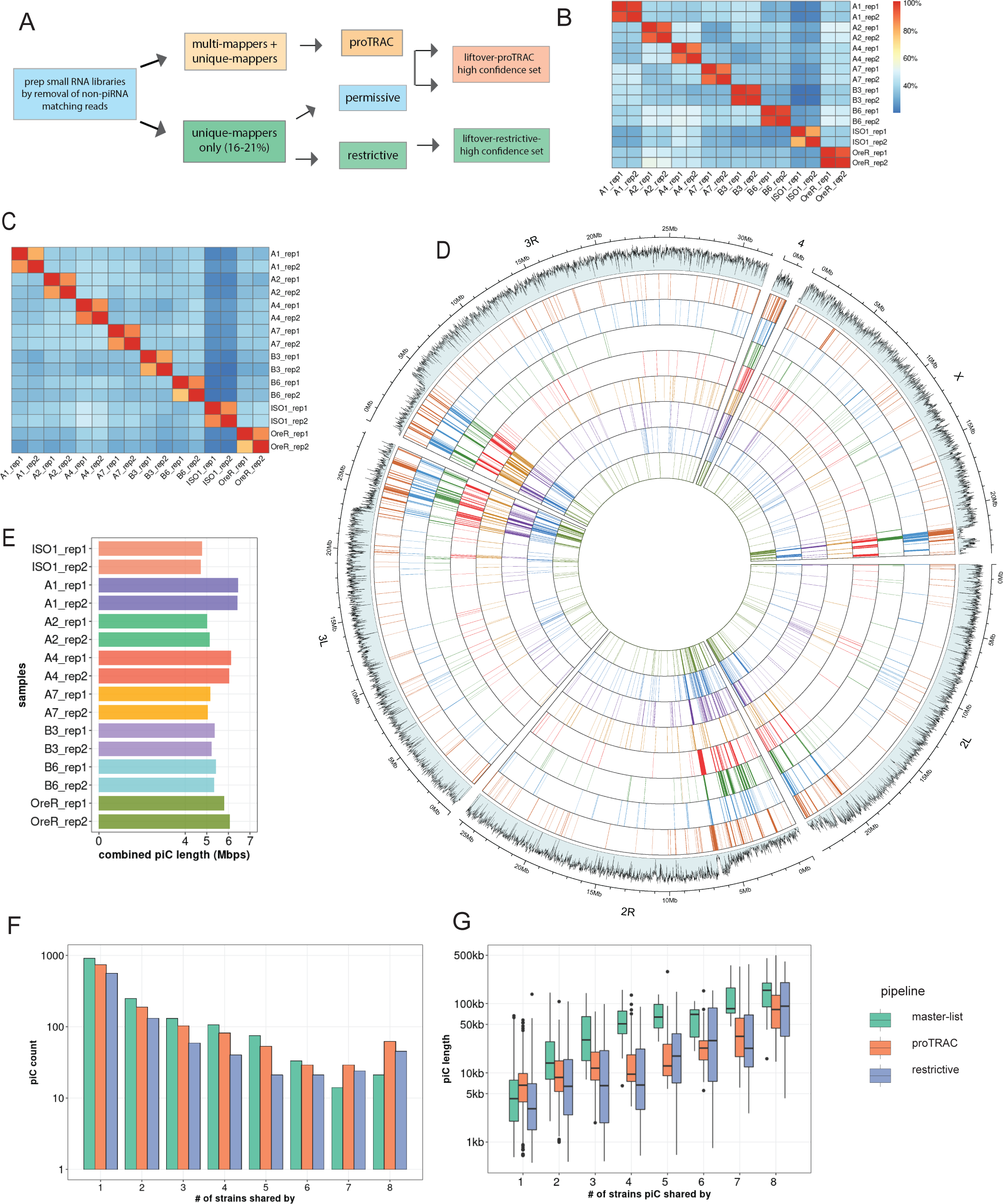
Inter-strain variability of piCs in *D. melanogaster* strains. (*A*) Summary of piC prediction and annotation pipeline using restrictive, permissive and proTRAC pipelines. *(B)* Cross-strain overlap (% of total piCs count) of independently predicted piCs for each replicate using the restrictive method. *(C)* Combined genomic piC size predicted from each replicate small RNA library independently in respective genome assemblies. *(D)* Genome-wide distribution of lifted-over piCs in 7 DSPR strains and reference iso-1 strain. Bars along the circumference represent presence of piCs in 10 kb bins for each chromosome. The outermost bar plot is piRNA mappability scores, followed by iso-1 piCs, followed by piCs of 7 DSPR strains. *(E)* Combined piC length predicted independently for each strain per small RNA library. *(F)* Population frequency of piCs in 7 DSPR strains and the reference iso-1 strain quantified after liftover to reference genome. *(G)* piC length distribution by population frequency in kilobase-pairs (kb) quantified after liftover to reference genome.

The piC predictions of the *restrictive* and *proTRAC* methods were both tested for reproducibility by comparing coordinates of piCs predicted independently from two replicates of each strain using bedtools. To quantify inter-replicate reproducibility in piC annotations, bedtools *intersect* was used with minimum required overlap of 20% of piC length for intra-strain replicates. Both *restrictive* and *proTRAC* methods yielded highly reproducible piC coordinates across replicates of each strain with >80% of piCs between two replicates overlapping over >75% of their respective length **(Fig. 1B,C, Supplementary Fig. 3A,B)**. However, inter-strain pairwise comparisons revealed that pairs of strains share only an average of ∼40% of their total piCs. The high confidence set of piCs from *proTRAC* that either exhibited high expression or were supported by uniquely mapping piRNAs, along with all piCs from the *restrictive* method for each replicate were combined to create a replicate-specific ‘master list’ of piCs.

Next, we used the master list of each strain to compare piC landscape across strains. To do so, we lifted over the piC coordinates from their own genome assembly to the iso-1 reference genome using the NCBI remapping tool (NCBI). We found that even with relaxed mapping criteria (0.33X to 3X coverage and >70% identity) to the reference genome, only ∼85-90% of all piCs from any given strain could be lifted over to the reference genome. Further inspection of piCs that failed lift-over revealed that they were relatively small clusters (500 - 5000 bp) and entirely absent in the reference genome or were large clusters (25-200 kb) that had undergone extensive structural rearrangements and therefore could not be lifted over to the reference genome using the NCBI remapping tool. To recover piCs that were >25 kb in size, apparently active in multiple strains but highly structurally variable (like *20A, 42AB* etc.), we manually identified and curated their coordinates by searching for the nearest annotated gene flanking the piCs in their respective genome assemblies (**Supplementary Fig. 2B**). We combined the results of the two prediction methods from two replicates to produce a collapsed master list of the piCs for each strain.

Genome-wide visualization of piC annotations across chromosomes reveals striking variability in piC landscape across the 8 strains **(Fig. 1D)**. In aggregate, the total amount of genomic DNA covered by active piCs in each strain ranged from 4.8 Mb to 6.3 Mb **(Fig. 1E)**. While their piC landscape is broadly similar in terms of being denser within peri-centromeric and telomeric heterochromatic regions (which are also characterized by low mappability scores) compared to euchromatic regions, it is readily apparent that many individual clusters are present in only one or a few strains, even within these heterochromatic regions. Smaller, euchromatic piCs are even more strikingly variable across strains despite being characterized by higher mappability scores **(Fig. 1F)**. Thus, from this first broad-scale view, it appears that the total amount of genomic space occupied by piC within each strain is largely similar, but the positions of piCs is highly variable across strains.

### Abundant strain-specific and strain-biased piCs

To quantitatively assess the frequency of piCs across the 8 strains, we quantified the overlap of piCs predicted independently for each of the 8 strains using the master list coordinates. To account for changes in size of piCs among strains, we required a minimum positional overlap of only 1 bp for piCs to be considered shared between strains. Even when using this non-conservative criterion, we found that 568 (*restrictive*) to 906 (*proTRAC*) of piCs are active only in a single or a few strains confirming that each strain has a very unique piC landscape **(Fig. 1F)**. The results are similar whether we used the piC predictions of the *restrictive* and *proTRAC* methods separately or the combined master list **(Supplementary Table 2)**. All strains exhibit 35-60 piCs that are strictly unique to that strain and another ∼30 piCs that could not be lifted-over and therefore are likely to be strain-specific also **(Supplementary Fig. 4B)**. Thus, we can conservatively estimate that each strain possesses between 50 and 100 piCs that are not shared by any of the other 7 strains examined. This is a conservative estimate because we only require 1 bp of overlap to consider piCs to be shared, so we may be overestimating the number of shared piCs. In addition, 142 and 69 piCs (‘*restrictive’* pipeline) are shared between two and three strains, respectively. All such piCs, shared by 3 strains or less are together termed as ‘rare’ piCs. Rare piCs are not only extremely abundant but also exhibit significant piRNA expression ranging from 20-100 RPM, which is comparable to previously described canonical piCs like *38C*, *80EF* and *traffic jam* 3’UTR **(Supplementary Table 3)**. Additionally, despite their small size, in aggregate, strain-specific piCs contribute a substantial portion of the total genomic span of piCs (average of ∼1 Mb) and 15-20% of the total piC genomic length of each strain **(Supplementary Fig. 4C)**.

Next, we examined the relationship between the size of piC and their level of sharing across strains. First, we note that median piC length predicted from each library is highly similar, with median length ranging from 5.7 kb to 7.5 kb **(Supplementary Fig. 4A)**. We find that piC size is positively correlated with the level of sharing across strains, and this correlation holds true for all prediction methods (Pearson *r* = 0.56 for proTRAC, 0.58 for restrictive, and 0.76 for master-list, *p-value* < 2.2e-16) (**Fig. 1G)**. In other words, piCs detected in a single or a minority of the strains (rare piCs) tend to be smaller (2-10 kb) than those shared by the majority of the strains (common piCs). If we posit that rare piCs represent evolutionarily younger piCs than common piCs, this relationship suggests that piCs are born relatively small and increase in size as they get older. Alternatively, larger piCs may be more evolutionarily stable than smaller ones. We note, however, that even large piCs can still be variable in activity across strains. For example, large well-known piCs like *42AB* and *38C* are still only active in 6 to 7 of the 8 strains (see below). Taken together, these results suggest that ovarian piCs are extremely labile and poorly conserved in activity across *D. melanogaster* strains.

### Extensive variability in piRNA expression of piCs

To illustrate the differences in activity of piCs among the 8 strains, we examine the piRNA coverage profiles for *42AB* and *82E*, two dual-stranded piCs **(Fig 2A,B)**. The *42AB* cluster has been extensively documented for its high piRNA expression (Brennecke et al. 2007; Klattenhoff et al. 2009). We present normalized coverage of uniquely mapping piRNAs to the respective *42AB* assemblies for the 8 strains using annotated flanking genes *Pld* and *jing* from both small RNA library replicates. Additionally, to examine differences in read coverage due to mappability, theoretical mappability scores are visualized along the length of the cluster in 100 bp bins **(Fig. 2A)**. Strains A1 and A7 have severely reduced (>20-fold) piRNA expression levels throughout *42AB* compared to the other strains. Similarly, *38C* – a highly productive dual-stranded piC in iso-1, exhibits significant variability in uniquely-mapping piRNAs across strains **(Supplementary Fig. S6)**. Since *42AB* and 38C are active in 6 out of 8 strains, it is most parsimonious to conclude that these piCs are relatively old but have lost activity in a subset of strains.

**Figure 2.**
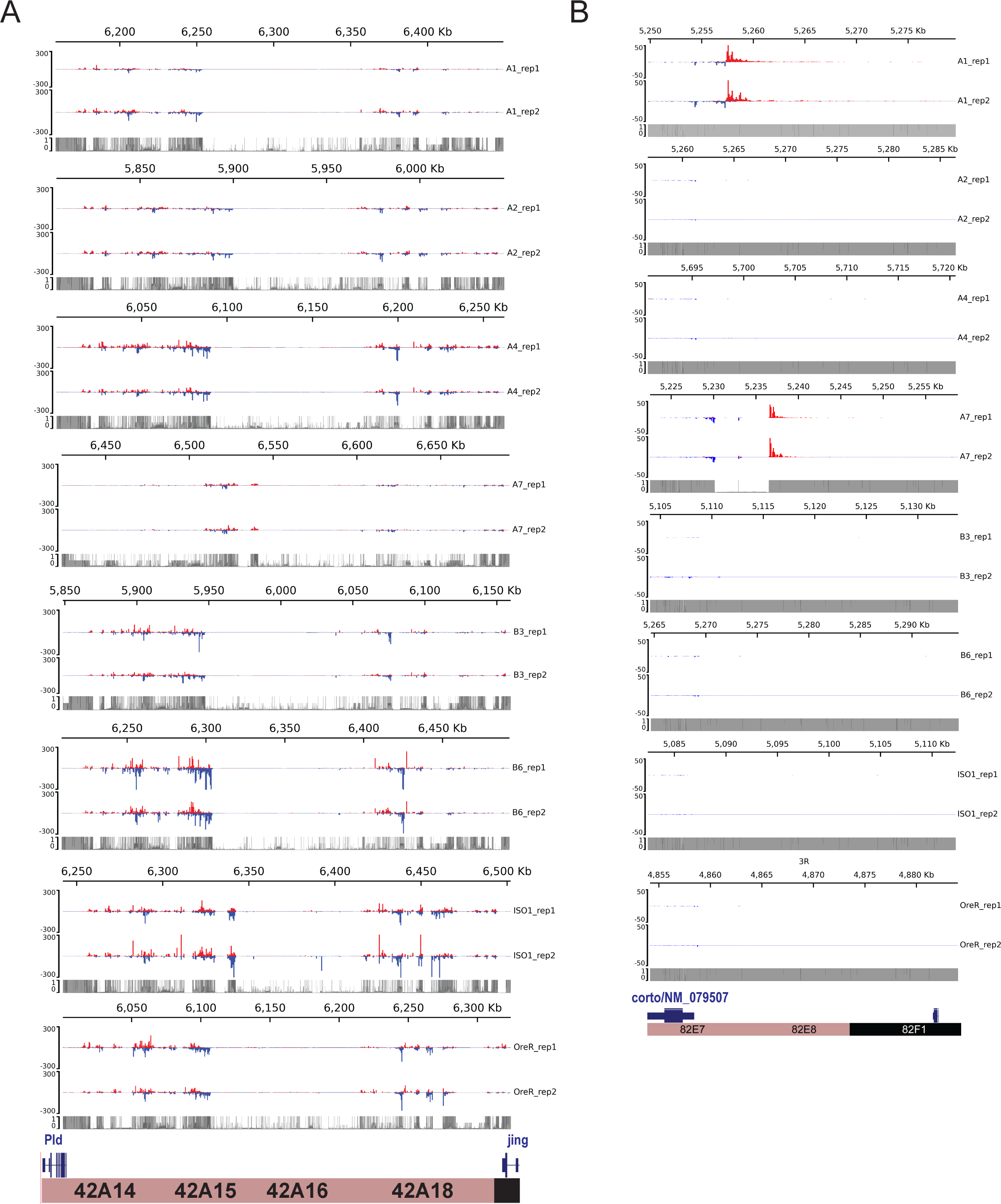
Natural variation in expression of uniquely mapping piRNAs from *42AB* and *82EF*. *(A)* Uniquely mapping piRNA expression profiles of *42AB* piCs for the 8 strains with two small RNA library replicates. Expression values are in reads per million (RPM) for 100 bp bins. Mappability scores (0-1) is shown for 100 bp bins of each respective *42AB* genomic assembly. *(B)* Uniquely mapping piRNA expression profile of *82EF* piCs for the 8 strains with two small RNA library replicates. Expression values are in reads per million (RPM) for 100 bp bins. Mappability scores (0-1) is shown for 100 bp bins of each respective *42AB* genomic assembly.

The *82E* cluster is a smaller piC (∼25 kb) we selected because it is also highly expressed in some strains but is euchromatic, overlapping the 5’ UTR of the *corto* gene. It is only identified as a piC in strains A1 and A7 in this study using the *restrictive* pipeline and piC expression profiles clearly show that piRNA expression across the *82E* region is only detectable in these two strains **(Fig. 2B)**. Other the other hand, *82E* has no detectable expression in 6 of 8 strains. Additionally, we show the average of normalized piRNA coverage from two replicates for the 8 strains in respective genome assemblies using the DrosOmics browser (Coronado-Zamora et al. 2023). Examples of variability in normalized piRNA expression across strains for piCs is shown. *80EF* (left) and *80F9* (right) are two peri-centromeric piCs on chr3L (**Supplementary Fig. 5A)**. *80EF*, a previously described Rhino-dependent piC, is a common piC, detected across all strains with significant piRNA production. *80F9*, however, is a less common piC with extremely variable piRNA coverage (50-fold) between strains and is annotated as a piC in only 6 strains. piRNA coverage of *Trypsin* genes-associated piC (left) and *eEF1alpha1* associated piC (right) in **Supplementary Fig. 5B** also are consistent with their detection by annotation pipelines in the respective strains. Comparison of such syntenic piCs between strains in their native genome assemblies provide validation of the variable activity of piCs across strains presented earlier from annotation pipelines **(Fig. 1G)**.

### Structural variation in piCs supports both ‘trap*’* and ‘*de novo’* models

To better understand the mutational processes underlying the divergence of piCs among strains, we examined the contribution of inter-strain structural variants (SVs), namely insertions and deletions (INDELs), within the piC genomic regions. INDELs are predicted to affect piCs differently under the *‘*trap*’* and *‘de novo’* models of piC evolution. Under the ‘trap’ model, we expect that the rate of insertions, likely representing the ‘trapping’ of TE insertions within common piCs would be balanced by that of deletions as to stabilize the size of the piCs. Thus, we expect a similar frequency of insertions and deletions within large piCs acting as traps. By contrast, we predict that clusters born ‘de novo’ would be dominated by insertions, likely corresponding to recent transposition events. To systematically detect SVs genome-wide for each of the strains relative to the iso-1 reference strain (Chakraborty et al. 2019; Solares et al. 2018), we mapped raw long sequencing reads for each strain to the iso-1 genome and called SVs using three independent SV callers (See Methods). SVs were then genotyped, filtered, and retained for downstream analyses if supported by at least two callers. SVs from all strains were then collapsed to construct a list of unique SVs that consisted of 2274 insertions and 4409 deletions relative to the reference genome. INDELs were then evolutionarily polarized into insertions vs. deletions by comparison of each variant to *D. simulans* and *D. sechellia* reference strains, which enable inference of the ancestral state (see methods). Polarization led to loss of ∼55% of INDEL calls as the ancestral or derived state of the loci could not be determined due to conflicts in calls between the two outgroup species. After this filtering, 1183 insertions and 1873 deletions were retained for analysis, of which 30% of insertions and 20% of deletions overlapped with master-list piCs **(Fig. 3A)**.

**Figure 3.**
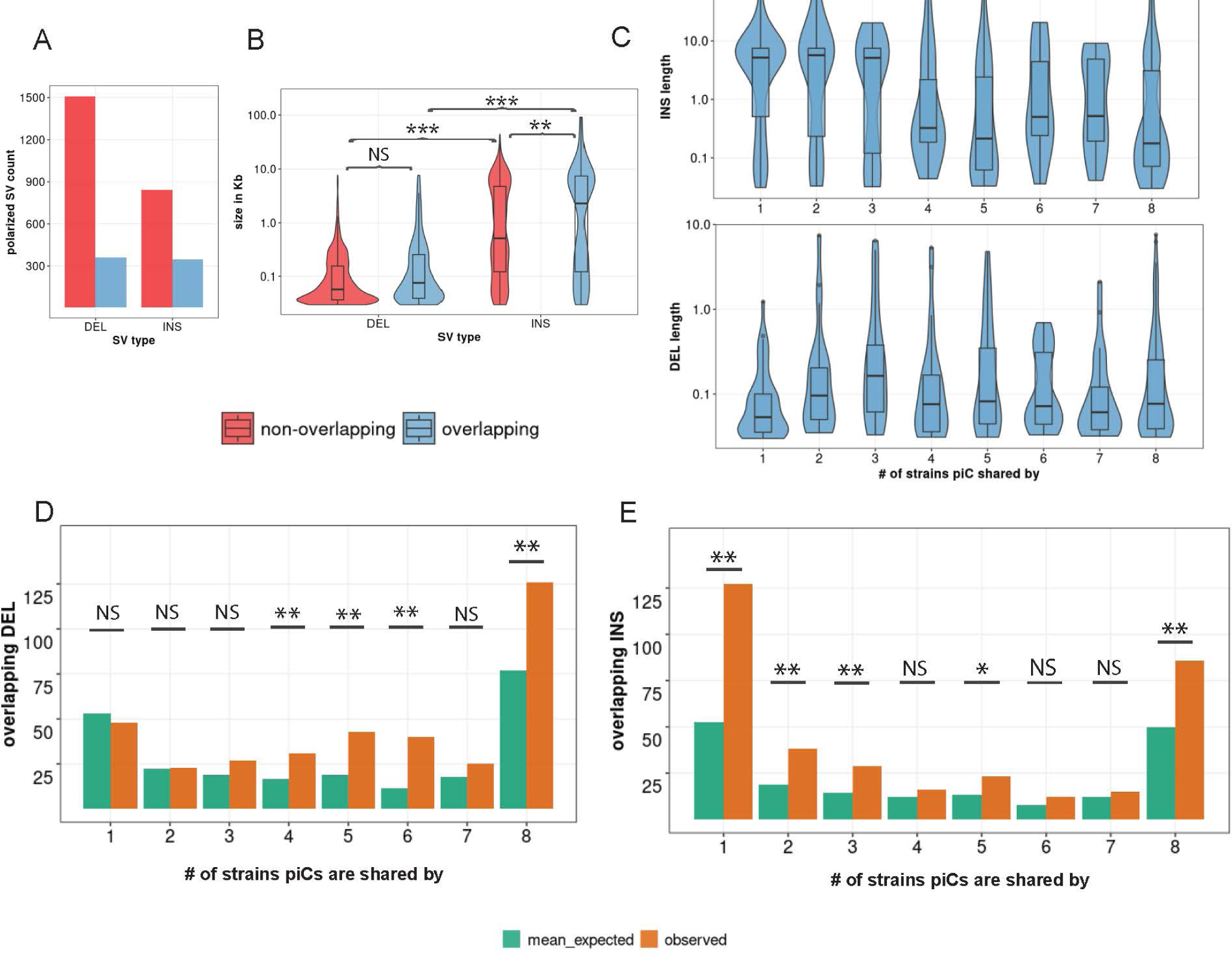
Common piCs exhibit ‘trap’ like sequence turnover. *(A)* Observed counts of INDELs overlapping and non-overlapping with piCs. *(B)* Length distribution of INDELs overlapping and non-overlapping with piCs. Significant differences are shown from Kruskal-Wallis test comparisons. *(C)* Length distribution of INDELs overlapping piCs grouped by the number of strains they are shared by (i.e., population frequency). *(D&E)* Enrichment analyses of deletion (DEL) and insertions (INS) variants in piCs carried out using poverlap. Variants overlapping with piCs of differing population frequencies is compared to expected mean overlap counts genome wide. (p-value <0.05=*, <0.005=**)

We examined the size distribution of INDELs overlapping piC and non-piC regions of the genome to then test the predictions for INDELs associated with piC variation. First, genome-wide, insertions range from 30 bp to 91 kb with a median of 612 bp, while deletions range from 30 bp to 7.6 kb with a lower median of 208 bp compared to insertions, which is consistent with previous structural variant profiling of *D. melanogaster* strains (Dopman and Hartl 2007; Zichner et al. 2013; Huang et al. 2014). However, insertions overlapping piCs have a median length of 2.2 kb, whereas insertions non-overlapping piCs have a smaller median length of 512 bp (Kruskal-Wallis test, χ^2^=10.812, df=1, *p*-value = 0.001). Meanwhile, deletions overlapping piCs have a similar length distribution than those non-overlapping piCs (Kruskal-Wallis test, χ^2^=0.72404, df= 1, *p*-value = 0.39) **(Fig. 3B)**. We also compared the length distribution of INDELs in piCs grouped by the number of strains with which they are shared. We found that strain-specific or rare piCs (shared by less than 4 strains) are associated with relatively large insertions (median length of 5.2 kb), whereas common piCs (shared by more than half of the strains) have a median insertion length of less than 1 kb **(Fig. 3C)**. However, the length distribution of deletions was similar between rare and common piCs. In sum, rare piCs are uniquely associated with relatively large insertions, which is consistent with the idea that these piCs emerged *de novo* from recent TE insertions.

Next, we tested whether piCs are enriched for INDELs relative to the rest of the genome. To do this, we compared the INDEL counts overlapping piCs for each of the cluster frequency categories with those expected based on 1000 sets of randomly shuffled INDELs. We found that deletions were significantly enriched in common piCs, but not in rare piCs **(Fig. 3D)**. Insertions were strongly enriched both in rare and common piCs **(Fig. 3E)**. These results may be confounded by the location of piCs within constitutive heterochromatin, where the rate of SVs is generally high (Chakraborty et al. 2021; Montgomery et al. 1991). However, we found that only ∼28% of all piCs lie within constitutive heterochromatin boundaries of the reference genome assembly and INDELs were significantly enriched in piCs even when we compared them to heterochromatic regions **(Supplementary Fig. S7)** (Riddle et al. 2011).Thus, the SV enrichment we observe within common piCs is unlikely to be solely driven by their location within constitutive heterochromatin. Overall, we conclude that piCs are subject to a high rate of structural genomic change relative to the rest of the genome, which likely contributes to their rapid evolutionary turnover. Additionally, we found that common piCs are enriched for both insertions and deletions, which is consistent with these clusters evolving as ‘traps’. By contrast, rare piCs are only enriched for insertions, which supports the notion that these are generally young clusters born ‘*de novo*’ from recent TE insertions.

### TE re-annotation of each strain uncovers ∼3 Mb of unannotated TE DNA

While our analysis of SVs within piCs supports that these events are important drivers of piC evolution, it does not directly address the role of TE activity. To assess the contribution of TEs to the composition and changes in the activity of piCs across strains, we carried out *de novo* annotation of TEs in each of the strain genome assemblies. This was necessary because many TE consensus sequences present in the reference TE library for *D. melanogaster* (FlyBase release 2019_05) were discovered and curated more two decades ago using primarily the iso-1 and Oregon-R strains (Bartolomé et al. 2002; Kaminker et al. 2002; Bowen and McDonald 2001). However, recent advances in long-read sequencing technology have provided a means to obtain a more unbiased view of the repetitive landscape of *Drosophila* genomes, revealing novel TE families (Ellison and Cao 2020; Rech et al. 2022; Han et al. 2022). We developed a TE annotation pipeline based on RepeatModeler2 (for discovery) (Flynn et al. 2020; Smit 1999), RepeatMasker (for annotation) and additional tools to distinguish novel TEs from known TEs and curate a comprehensive TE library for the 8 strains used in this study **(Fig. 4A**, See methods).

**Figure 4.**
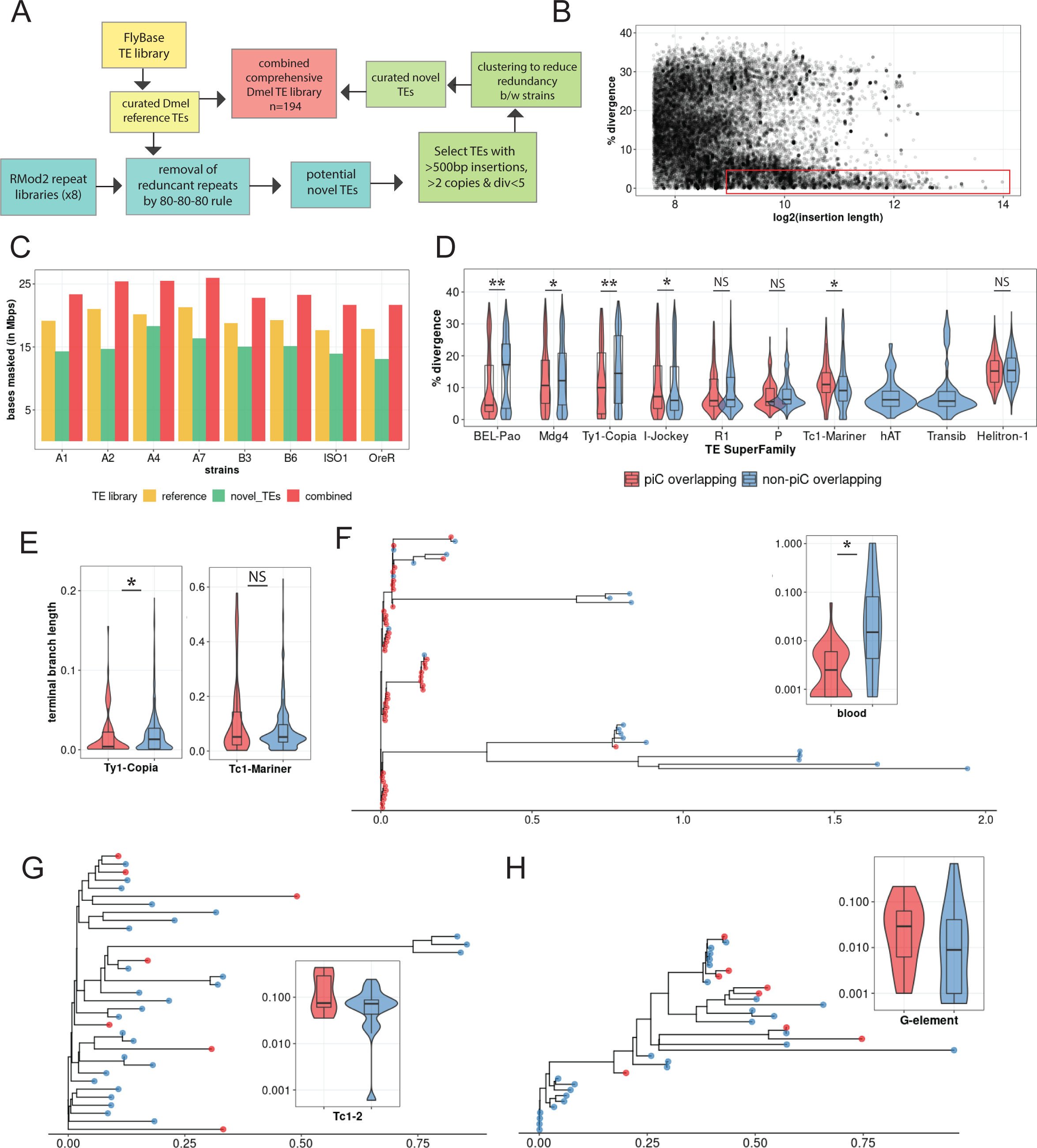
*de novo* TE annotation uncovers ∼3Mb of hidden TEs and reveals strong associations of young LTR TEs with piCs than any other TE subclasses. *(A)* TE annotation pipeline using RepeatModeler2 and RepeatMasker to create the comprehensive TE library. *(B)* Abundance of extremely similar and long TE insertions from RepeatMasker output of strain B6 using novel TE consensus library. *(C)* Differences in million base-pairs (Mbps) masked in RepeatMasker results using novel-only, reference-only, and combined TE library. *(D)* Divergence estimates for all defragmented iso-1 insertions (>250 bp) from RepeatMasker output. Insertions with >1 bp overlap with master-list iso-1 piCs were considered piC overlapping. Difference between groups is tested by Wilcox ranked-sum test. *(E)* Terminal branch length for all iso-1 insertions from *Ty1*/*Copia* and *Tc1*/*Mariner* superfamilies from maximum likelihood trees*. (F-H)* Maximum likelihood trees constructed from all defragmented insertions for *blood*, *Tc1-2*, and *G-element* families and the inset shows terminal branch length quantification. Difference between groups is tested by Wilcox ranked-sum test; p-value <0.05=*, <0.005= **, <0.0005=***.

TE family sequence assemblies by RepeatModeler2 for each strain were aligned to the reference TEs using the 80-80-80 rule (Wicker et al. 2007). RepeatModeler2 sequences already present in the reference TE library were removed. Next, the remaining RepeatModeler2 sequences were used to re-mask the genomes to examine them for novel TE families. Over 5000 insertions (>500 bp in size, median of 676 bp), very similar (<5% divergence) to their respective RepeatModeler2 family consensus are discovered for each strain (example of strain B6 in **Fig. 4B**). These novel insertions resulted in masking of an additional 2.5 Mb to 4 Mb in each genome assembly that would have been missed or mis-annotated as highly diverged insertions by masking with only the reference TE library **(Fig. 4C)**. In summary, a refined and comprehensive TE library was created with a combination of 129 reference TE consensus sequences and 45 uncharacterized consensus sequences that capture all TE insertions genome-wide and reflect their relative age.

### piC TE composition is represented by younger LTR insertions than any other TE subclass

Using the new TE library described above, we sought to compare the age and composition of TEs within piCs to that of the rest of the genome. To infer the age of each family, we use the median sequence divergence of each insertion to their family consensus. To examine TE composition, we grouped TEs into the major subclasses and superfamilies represented in *Drosophila*: non-LTR retrotransposons (LINE), LTR retrotransposons (*Ty1*/*copia*; *Ty3*/*mdg4*; *BEL*/*Pao* superfamilies), Rolling Circle (RC) transposons, and cut-and-paste DNA transposons. We found that TE copies from all three LTR superfamilies were significantly younger in piCs than non-piC regions **(Fig. 4D)**. Conversely, TE copies from LINE, RC and DNA subclasses were not significantly different in age in piCs than in non-piC regions. To corroborate these results using an independent method to date insertions, we built phylogenetic trees from all copies for one LTR superfamily (*Ty1*/*copia*) and one DNA transposon superfamily (*Tc1*/*mariner*) and used terminal branch lengths to estimate their relative age (Carr et al. 2012). We chose these superfamilies because they were of moderate abundance and therefore manageable for multiple sequence alignments and phylogenetic analyses. The results of these analyses yielded the same trend observed genome-wide using sequence divergence from consensus sequences whereby the *Ty1*/*copia* LTR retrotransposons (*n*=135) overlapping piCs are significantly younger than non-overlapping ones, while *Tc1*/*mariner* (*n*=89) DNA transposons show no such bias **(Fig. 4E)**.

To examine whether these trends hold at the level of individual TE families, we selected one family with moderate copy number from the LTR, LINE and DNA subclass and compared the age of piC-overlapping and non-overlapping copies within each family. As a representative *Ty3*/*mdg4* LTR superfamily, we analyzed *blood*, a family with 63 copies in the iso-1 strain that is known to be transpositionally active (Bingham and Chapman 1986; Kofler et al. 2015). Consistent with the trend observed at the level of the LTR superfamily, we found that 43 out of 63 *blood* insertions are associated with piCs. Most of these are very recent insertions with median terminal branch length of <0.002, which is significantly shorter than of insertions not overlapping piCs (Wilcoxon rank sum test, *p*-value = 0.014) **(Fig. 4F)**. In other words, piC overlapping *blood* insertions are significantly younger than the non-overlapping ones. As a representative of the *Tc1/mariner* superfamily of DNA transposons, we analyzed *Tc1*-2, a family with 35 copies in the iso-1 genome. Consistent with the trend observed at the level of the entire superfamily, the age of *Tc1-2* copies overlapping piC is not significantly different than that of non-piC overlapping copies (Wilcoxon rank sum test, *p*-value = 0.903) **(Fig. 4G)**. Analyzing the *G*-element LINE family, which counts 35 copies in iso-1 and is still active (di Nocera et al. 1986), we found that the age of piC-overlapping copies is not significantly different from non-overlapping copies (Wilcoxon rank sum test, *p*-value 0.855) and the youngest *G*-element insertions according to terminal branch length do not overlap piCs **(Fig. 4H)**. Taken together, these results suggest that young LTR retrotransposon insertions tend to be enriched in piCs, but this trend is not observed for other TE subclasses and superfamilies.

### A small subset of active LTR retrotransposon families give rise to young piCs

To examine the role of recent transposition events in driving piC sequence composition, we established a set of non-reference TE insertions (absent from the reference genome) in each of the 8 strains using the raw long read data available for each. Briefly, we applied TLDR (a long-read TE insertion detection tool) (Ewing et al. 2020) with a cut-off of at least 2 supporting reads per 10X genome coverage to remove false positives and enrich for germline insertions (see Methods). Using these parameters, we identified 285 to 857 non-reference TE insertions for each of the 7 DSPR lines but only 75 insertions for iso-1, which is expected since the reference genome is also derived from the iso-1 strain. Presumably, the 75 non-reference insertions for iso-1 reflect the use of different isolates for the reference genome assembly and for the long read data. Further clustering and parsing of all non-reference insertions across the 8 strains resulted in a list of 3545 unique TE insertions of at least 200 bp in length. These insertions belong to 165 of the 184 different families in our TE library. Ninety-four of these 165 TE families were classified as “active” when they included at least 5 non-reference insertions shared by no more than 2 strains, while the other 98 families were classified as “inactive” **(Fig. 5A)**.

**Figure 5.**
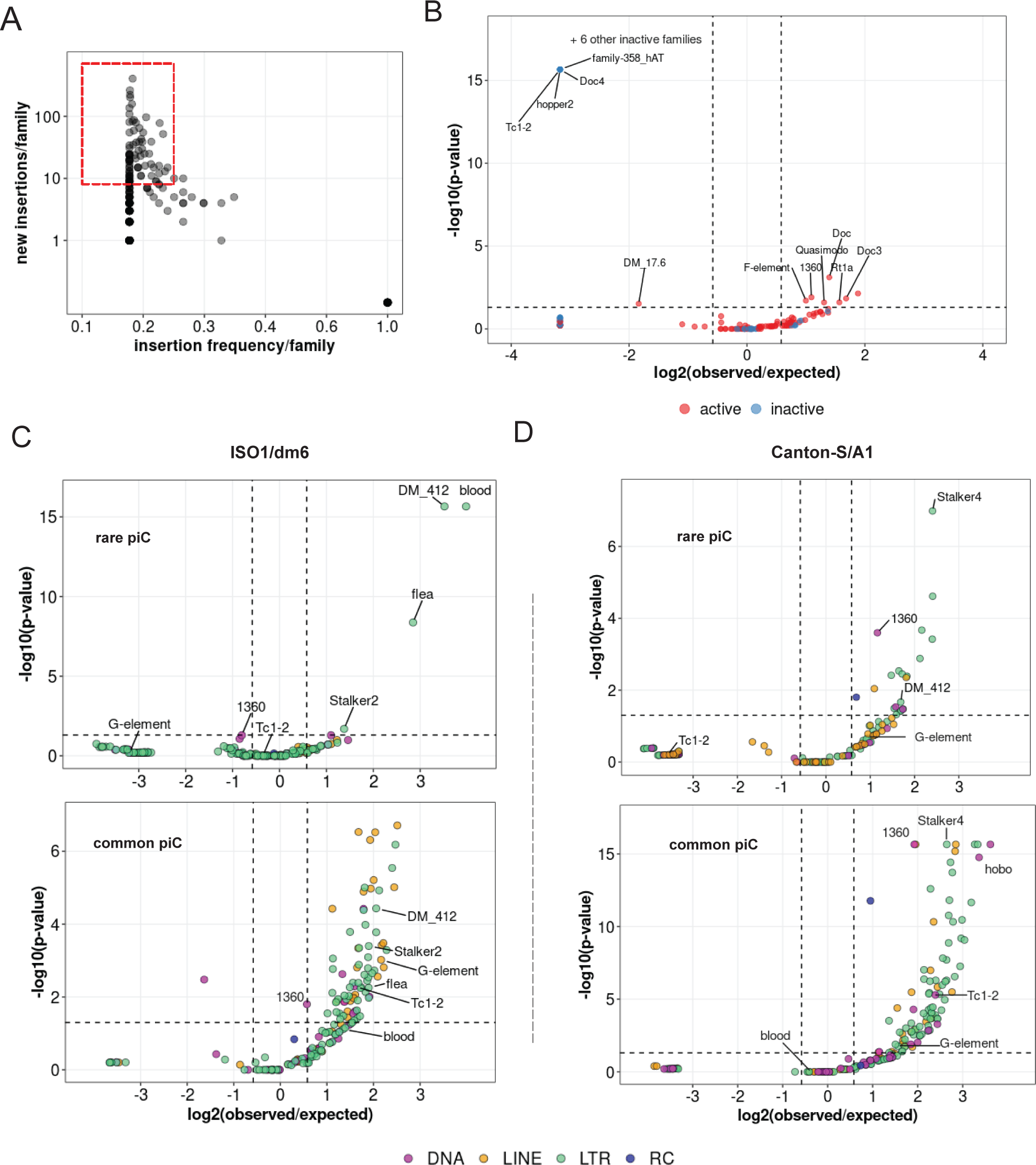
Insertions of only few active LTR families associates with rare piCs. *(A)* Scatter plot of non-reference TE insertion counts and mean population frequencies of 184 TE families. Red box highlights selected TEs classified as ‘active’.*(B)* Enrichment analyses of TE families in master-list piCs using random shuffling. Y-axis P-values are from binomial tests conducted to compare observed counts to expected average overlaps of de novo TE insertions to piCs for each family. *(C-D)* Enrichment analyses of TE families in master-list rare and common piCs of A1/Can-S strain using random shuffling.*P*-values on y-axes are from binomial test conducted to compare observed counts to expected average overlaps of de novo TE insertions to piCs for each family. Names of some of the statistically significant families are shown. *(E-F)* Enrichment analyses of TE families in the master-list of rare and common piCs of iso-1 using random huffling. *P*-values on y-axes are from binomial tests conducted to compare observed counts to expected verage overlaps of de novo TE insertions to piCs for each family. Names of some of the statistically ignificant families are shown.

We used this compendium of insertions to test whether TE families are significantly enriched or depleted within piCs based on their activity, using a binomial test to compare the observed overlaps with piCs to the average overlaps expected from 1000 random reshufflings of piC coordinates (see Methods) (Kapusta et al. 2013). This analysis revealed that 7 active TE families are significantly enriched in piCs, while 1 active and 10 inactive families are significantly depleted in piCs **(Fig. 5B)**. All the inactive TE families that are significantly depleted in piCs belong to either DNA or LINE subclasses, while the only active family depleted in piCs was 17.6, a *Ty1/copia* superfamily member (Inouye et al. 1986) with 79 non-reference insertions. These results are consistent with the prediction of the ‘trap’ of model that piCs are enriched for active TE families but are also composed of inactive families.

Next, we sought to distinguish family-level enrichment of TE insertions within rare ‘*de novo*’ piCs (smaller piCs (<10 kb) shared by no more than 3 strains) and within common larger ‘trap’-like piCs (>10 kb, shared by at least 5 strains). To increase statistical power for this analysis, we used all TE sequences annotated by RepeatMasker in each genome, instead of only non-reference insertions. For iso-1, we found that only 4 TE families are significantly enriched within young piCs. Interestingly, all 4 are LTR retrotransposon families of the *Ty3/Mdg4* superfamily (*blood, 412, flea, Stalker-2*) and all except *flea* belong to the *Mdg1* lineage (Bertocchi et al. 2020; Costas et al. 2001), suggesting that this lineage of elements may be prone to seed new piCs. All four families are also classified as active in this study as well as previous studies that examined TE insertion frequency among *D. melanogaster* populations (Kofler et al. 2015; Kelleher and Barbash 2013). By contrast, we found that numerous active and inactive families from all TE subclasses are significantly enriched in large and common “trap-like” piCs, largely representative of the overall TE landscape of *D. melanogaster* **(Fig. 5C)**. In the A1 genome, 20 TE families are significantly enriched in rare piCs. Again, these are predominantly LTR retrotransposons (14 families), but 4 DNA transposon families and 2 LINE families are also significantly enriched. **(Fig. 5D)**. Interestingly, *blood* insertions are neither enriched nor depleted in common piCs of both strains. Taken together, these analyses yield a contrasting portrait of TE composition in the two major types of piCs.

### TE composition of piCs captures distinct steps in piC evolution

To further illuminate the evolution of piCs, we analyzed how the overall age and distribution of TEs of piCs change as they become more frequent, and presumably older. We plotted mean percent divergence of individual TE insertions to their consensus sequences (a measure of TE age) across piCs and their flanking non-piC regions for each piC frequency class (strain-specific or shared by 2-8 strains). We found that the divergence of TE insertions in rare piCs (shared by 3 or less strains) is markedly lower (3.5-5%) than in their flanking regions (10-15%) **(Fig. 6A)**. In addition, the divergence of TE copies within piCs increases gradually with the frequency of the piCs to the extent that for the most common piCs (shared by 7 and 8 strains) the mean percent TE divergence is only slightly lower than in their flanking genomic regions. This apparent increase in age of TE insertions as piCs become more frequent provides weight to the inference that more common piCs represent evolutionary older clusters relative to those that are strain-specific or rare. It also suggests that piCs are born from *de novo* TE insertions and grow by gradual accretion of TEs over time.

**Figure 6.**
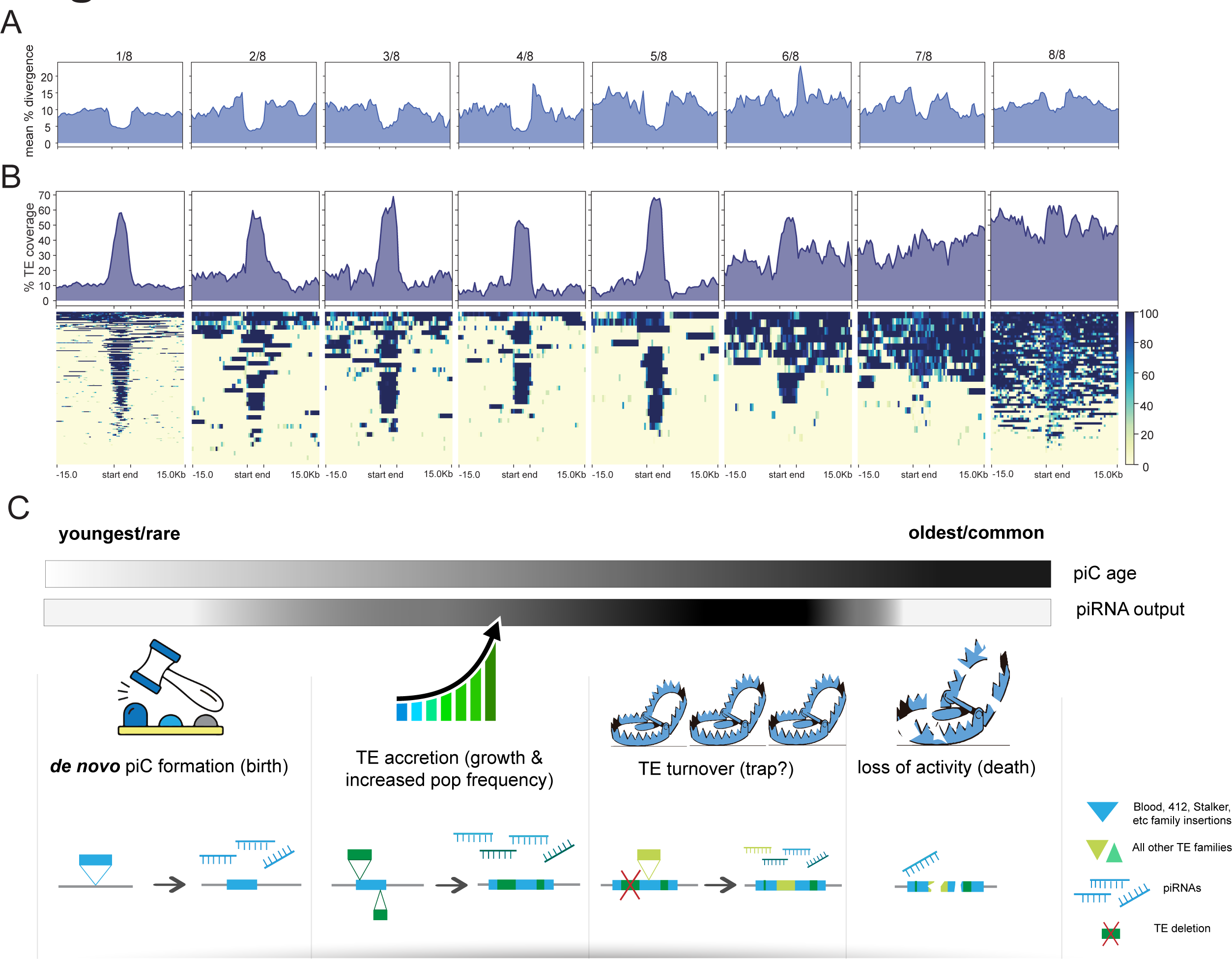
Age and distribution of TEs provide a portrait of intra-specific piC evolution. *(A)* Summary profile plot of ercent divergence in 500 bp windows in scaled piC regions and flanking non-piC regions of the reference e. Each group represents piCs that are shared by 1/8 to 8/8 strains.*(B)* Summary profile plot and heatmap of E coverage in 500 bp windows in scaled piC regions and flanking −/+ 15 kb non-piC regions of the reference e.*(C)* New unified model of piC evolution in four steps from left to right.

To further test this idea, we examined how TE coverage within piCs and surrounding regions change as piCs become more common and presumably older **(Fig. 6B)**. First, we observed that piCs generally exhibit significantly higher TE coverage than their flanking genomic regions. Second, we found that strain-specific piCs, which presumably represent the youngest piCs, exhibits high mean TE coverage in the middle at >60% (on average 60 out every 100 bp is comprised of TE sequence), which drops dramatically at the edges of piC coordinates to <20% **(Fig. 6B)**. In contrast, more common piC groups exhibit consistently higher TE coverage across their entire length. This pattern is consistent with a birth and growth process where a piC emerges from individual TE insertion, but piRNA production spreads to flanking TE insertions as they insert near or within the piC.

## Discussion

To study piC evolution at fine-scale resolution in *D. melanogaster*, we used population genomics methods to characterize piC variation across eight inbred strains. A crucial asset was the availability of high-quality genome assemblies for these strains (Chakraborty et al. 2019). This enabled us to produce *de novo* annotation of piCs for each strain after mapping inferred piRNAs from ovarian small RNA libraries we constructed and sequenced for two biological replicates sampled six months apart. Our piC annotations for the two replicates exhibited high reproducibility with >95% of piCs annotated in one replicate found in the second replicate **(Fig. 1A,B, Supplementary Fig. 3)**. Also, to understand variation in sequence composition and age of piCs, it was necessary to produce libraries of TE consensus sequences representative of the eight strains analyzed here. By performing *de novo* discovery and re-annotation of TE families for each genome, we identified 49 novel TE consensus **(Fig. 4)**. While further investigation is required to examine their evolutionary origins and relationship to known TE families, it appears that many of the novel TE families we annotated were highly diverged from known families and often “hidden” in highly repetitive regions that would likely be poorly assembled in short-read genome assemblies. These results stress the benefits of high-quality genome assemblies and the necessity to perform *de novo* TE discovery when new individuals, strains or geographical isolates are considered. This is true even for model species like *D. melanogaster,* where TEs have been extensively cataloged, because previous TE identifications were mostly based on a single reference genome. Robust annotation of piCs and TEs allowed us to compare in detail the activity and TE composition of piCs across strains and test the generality of two contrasting models of piC evolution.

We sought to distinguish which of the two models – ‘*de novo*’ or ‘trap’ -- best captures piC evolution in *D. melanogaster*. First, the chromosomal location and distribution of piCs sampled in this study are largely consistent with the ‘*de novo*’ model. We do, however, identify a small subset of piCs (20-30) that are shared across most or all strains, a characteristic that is congruent with the ‘trap’ model **(Fig. 1C,E)**. Second, we observe extensive variation in piC activity and abundant strain-specific piCs, features supporting the ‘*de novo*’ model **(Fig. 1F)**. We also find a positive correlation between piC length and frequency suggesting that piCs are born small but grow in size as they become more common in the population **(Fig. 1G)**. Third, INDELs associated with piCs exhibit predicted signatures of both ‘*de novo*’ and ‘*trap*’ models but only in specific groups of piCs – rare and common respectively **(Fig. 3)**. Fourth, the age of TE insertions within piCs are consistent with the ‘trap’ model, whereby active or recently active TE families are enriched, while inactive ones are depleted **(Fig. 4,5)**. Thus, our findings recapitulate predictions of both ‘*de novo*’ and trap models of piC evolution.

Overall, our results support the idea that piCs are primarily born ‘*de novo*’, but a small subset of large heterochromatic clusters are more evolutionarily stable and appear to behave as ‘traps’. For example, piCs like *flamenco*, *20A*, and *h52-3* show robust piRNA expression in all strains analyzed. However, several large piCs like *42AB, 38C, Myo81F* show significant loss in piRNA production in one or multiple strains **(Fig. 2, Supplementary Fig 6)**. We were able to rule out mappability differences as a confounding factor **(Fig. 2)**. What then causes the loss in piRNA production from large heterochromatic piCs? Our current lack of understanding of the cis-regulatory requirements for piC activity makes it difficult to determine whether changes in piRNA production are caused by genetic or epigenetic changes in the piCs. We and others (Wierzbicki et al. 2021b; Ellison and Cao 2020) observe considerable structural variation among strains in large peri-centromeric piCs, including *38C* and *42AB.* It is possible that such structural changes result in changes in piRNA production, but further studies are needed to elucidate the mechanisms by which large and seemingly stable piCs lose their activity.

What can TE composition of piCs tell us about the coevolution of TEs and piCs? As predicted and previously reported (Chen et al. 2021; Brennecke et al. 2007; Ellison and Cao 2020; Wierzbicki et al. 2023; Kofler et al. 2015), we found that diverse TE families (from all subclasses) are enriched in large, common piCs **(Fig. 5)**. However, we found that this trend is driven mostly by younger LTR insertions **(Fig. 4)**. This enrichment may be explained by selection against *de novo* TE insertions in gene-rich euchromatic regions, which leads to unrestricted accumulation of TEs in heterochromatic regions and where purifying selection is also weak (Blumenstiel et al. 2002; Schrider et al. 2013; Charlesworth and Langley 1989; Dolgin and Charlesworth 2006). However, our genome-wide analysis revealed that SV enrichment in common piCs cannot be completely explained by their overlap with heterochromatin **(Supplementary Fig 7)**. Thus, the enrichment of LTR elements within large common piCs may be driven in part by their insertion preference and/or by selection for their repression.

Are particular TEs prone to give rise to piCs *de novo*? To answer this question, we tested for enrichment of individual TE families within rare piCs. Interestingly, only a small set of retrotransposon families are enriched within such clusters and most belong to the *mdg1* subclade of LTR retrotransposons such as *blood*, *412,* and *Stalker* families **(Fig. 5C,D)** (Kapitonov and Jurka 2003; Nefedova and Kim 2009). Why would these elements be prone to seed new piCs? We hypothesize that this may be linked to their propensity to produce double-stranded RNA (dsRNA) and endogenous siRNAs. It is well known that many LTR and non-LTR retrotransposons possess bi-directional promoters that can result in the formation of dsRNAs that stimulate the production of siRNAs (Hung and Slotkin 2021; Watanabe et al. 2008), including *blood* and *412* (Russo et al. 2016). Because endo-siRNAs production has been associated with the formation of transgenic piCs in flies (Olovnikov et al. 2013; Le Thomas et al. 2014), it is possible that the propensity of these retrotransposon families to produce dsRNAs nucleate the formation of piCs. This idea has received support in recent study showing that endogenous siRNA production precedes piRNA cluster formation and maternal inheritance of these siRNAs is required for licensing of piRNA clusters (Luo et al. 2022). Thus, we hypothesize that a subset of retrotransposon insertions prone to produce dsRNA enter the endo-siRNA pathway which in turn promote the birth of new piCs at these loci. In other words, the rapid evolution of piCs among *D. melanogaster* strains may be driven by the activity of a few TE families.

Based on all these observations, we propose a ’birth-and-death’ model of piC evolution, which combines components of both ‘trap’ and ‘*de novo’* models (Moon et al. 2018; Shpiz et al. 2014; Zhang et al. 2020; Bergman et al. 2006). In this model **(Fig 6C)**, we posit that piCs form frequently throughout the genome, mostly from recent TE insertions, with certain LTR retrotransposon families making a stronger contribution to seeding new piCs. Newly emerged piCs may increase in frequency and size due to natural selection or drift, depending on factors such as their propensity to trigger genomic autoimmunity (Blumenstiel et al. 2016; Lee and Karpen 2017), ectopic recombination (Petrov et al. 2003; Sentmanat and Elgin 2012) and the establishment of a chromatin environment conducive for piRNA production (le Thomas et al. 2014). Over time, these stabilized clusters may grow by ‘trapping’ additional TE insertions, which will eventually result in large heterochromatin clusters such as *flamenco* and *42AB*. Due to the host’s limited capacity to maintain such piCs without incurring a fitness cost, those clusters that lack piRNAs targeting active TEs may gradually lose activity or become dispensable (Gebert et al. 2021). Further studies are warranted to test this “birth-and-death” model. Our study provides a first in-depth view of piC evolution in *D. melanogaster* that is likely to stimulate other comparative studies of piRNA evolution.

## Materials and Methods

### Fly stocks

DSPR founder stocks of A1 (b1_paired), A2 (b3841_paired), A4 (b1_3852), A7 (t7_paired), B3 (b3864_paired), B6 (t1_paired) and Oregon-R were a gift from Anthony Long (UCI). iso-1 reference strain (#2057) was obtained from Bloomington Drosophila Stock Center. All stocks were maintained on standard cornmeal medium at 22°C under a 12-hr day/night cycle.

### Small RNA library construction and sequencing

Small RNA libraries were constructed by size fractionation on urea-polyacrylamide gel electrophoresis as described in Ma *et al*. 2021. Libraries were constructed for two biological replicates per strain, from ovaries collected at times separated by 25 weeks. Briefly, ovaries were dissected from 25-30 yeast-fed adult females of 4-6 day old and total RNA was extracted using TRIzol reagent and quantified with NanoDrop. Small RNAs of 17-29 nt length were size fractionated from 10 µg of total RNA on Novex TBE-Urea Gels, 15% (Thermo Fischer EC6885BOX) using ZR small-RNA ladder (ZymoResearch, R1090) as reference. Small RNAs in excised gel fragments were first eluted in 500µL 0.3M NaCl and kept on an agitator for 16 hrs. Small RNAs were then precipitated with 2 volumes of chilled isopropanol and 1 µL of 20 mg/mL glycogen. Small RNA pellet was then washed with chilled 75% ethanol and eluted in 10µL of freshly made 50% (w/v) PEG-8000 to enhance 3′ end ligation efficiency. Library preparation was carried out using half of eluted small RNAs (5 µL) for each replicate with NEB Small RNA library preparation kit (E7300) as per manufacturer’s protocol. All libraries were PCR amplified to 14 cycles, visualized on a 2% agarose gel, and purified with NEB Monarch PCR & DNA Cleanup Kit (T1030S). All libraries were quantified using Qubit 3.0, pooled into replicate-1 and replicate-2 groups, and analyzed on Agilent Bioanalyzer. Single end 75 bp Illumina sequencing was carried out for all libraries on NextSeq500 at Cornell Biotechnology Resource Center and few select libraries were re-sequenced if a minimum of 10 million reads was not obtained in first round.

### Processing of small RNA libraries

Reads were first trimmed of the adapter sequences and quality filtered using cutadapt (v3.4)(Martin 2011). Read length distribution, per sequence quality and duplication level was obtained from FastQC (Andrews and others 2010). Reads mapping to annotated miRNAs (Kozomara et al. 2019), other non-coding RNAs like rRNA, snRNA, tRNA, and snoRNA sequences (Hoskins et al. 2015) were removed using bowtie (-v 2-k 1-y –un-S) and a combined custom reference of non-coding small RNAs. The remaining 23-29 nt genome-mapping reads were retained as piRNA reads.

### piC annotation

Active piRNA cluster (piC) annotation was conducted independently for each replicate of each strain using a custom pipeline adapted from previously described methods (Mohn et al. 2014; Rosenkranz and Zischler 2012). Genome mapping 23-29 reads, filtered from annotated non-piRNA small RNA genes, were mapped to respective genome assemblies using bowtie -n 1 −l 12 -a -m 1 -y -S and resulting unique alignments were separated from unmapped reads using samtools in bam files (Langmead et al. 2009; Li et al. 2009). bedtools *makewindows* (Quinlan and Hall 2010) was used to create 500 bp bookended windows from 7 DSPR genome assemblies (Chakraborty et al. 2019) and iso-1 reference assembly (Hoskins et al. 2015) followed by bedtools *coverage* to calculate uniquely mapped piRNA reads per million (RPM) per window from bam files. Windows with piRNA expression of 2 RPM or more were merged if located within 10 kb of each other into piRNA expression domains. RPKM values for piRNA expression was calculated for such domains (ranged from 500 bp to 330 kb).

piC annotation from merged domains was conducted in two modes - permissive and restrictive. First piRNA reads that uniquely map to selected domains were recovered and separated for each domain using samtools to quantify unique piRNA sequences per domain. Additionally, theoretical mappability scores of 0-1 for 25 nt reads was computed using GEM (Derrien et al. 2012) for bookended 500 bp windows for each genome assembly. For the *restrictive* mode of annotation - domains with at least 8 unique piRNAs per kb per million total piRNA reads were selected and then merged if interrupted by low mappability region of 10 kb or less. Similarly, for the permissive mode, domains with at least 2 piRNAs per kb per million were selected and then merged if interrupted by low mappability region of 15 kb or less. While the permissive mode had very relaxed parameters and likely produced many false positive predictions, the primary function of this mode was simply to provide unique piRNA support for the piRNAs detected from proTRAC method (see below), which utilizes multi-mapping piRNAs and predicts the majority of piRNAs in extremely low-mappability regions.

### Alternate *de novo* annotation of piCs by proTRAC

The same processed small RNA libraries as described above were used for alternative piC annotation using proTRAC-v2.4.4 (Rosenkranz and Zischler 2012). Specifically, for proTRAC analysis, each library was collapsed to include only unique piRNA sequences using TBr2_collapse of NGS-toolbox (Rosenkranz et al. 2015) and mapped to respective genomes using bowtie -n 1 - l 12-a --best --strata --quiet -y-chunkmbs 1024. proTRAC 2.4.4 was run on sam files generated from bowtie with the following parameters-swsize 500-swincr 100-clsize 500-1Tor10A 0.6-pimin 23-clhitsn 10-pdens 0.2-pti followed by removal of clusters with normalized multi-mapped piRNA coverage of 25 or less. proTRAC piC annotation was extracted from resulting clusters.gtf files.

### liftOver (remap) of piCs and genes for DSPR genome assemblies

piCs annotated from the above methods for each strain were lifted-over to the iso-1 reference genome (Release 6) using NCBI remap (NCBI Genome Remapping Service). Briefly piC coordinates from the custom *restrictive* pipeline and *proTRAC* were lifted-over from each strain to iso-1. Remap parameters chosen for identification of all piCs with minimum alignment coverage of 0.3 and maximum expansion or contraction of 3X, allowing for clusters with strain-specific structural variation to be detected. Similarly, gene annotation from the reference genome was also lifted over to the DSPR genomes but with higher mapping stringency of minimum alignment coverage of 0.9.

### Manual curation and collapsed replicate annotations

Recovery of complete piC regions primarily depends on expression and density of uniquely mapped piRNAs along the length of the cluster. Clusters identified independently from two different strains may differ in length due to natural variation in piRNA expression or density along the length of the cluster. To identify homologous clusters across strains despite changed boundaries due to structural variation in piCs, most heterochromatic piCs were examined in the IGV browser. Clusters resulting from merged bins across annotated protein-coding genes were unmerged into separate clusters. Any cluster partitioned into multiple smaller clusters due to lack of any uniquely mapping piRNAs for more than 20 kb were merged to recover the complete cluster. All strain genome assembly-match piC annotations from *restrictive*, *proTRAC* and *master-list* are provided in **Supplementary Table S2,** whereas curated and replicate collapsed annotations are provided in **Supplementary Table S3**.

### Structural variation detection and filtering

Raw long reads for the 7 DSPR strains, the iso-1 reference strain, *D. simulans* (wxd1) and *D. sechellia* (sech25) were mapped to the *D. melanogaster* iso-1 release 6 (GCA_000001215.4) without the Y chromosome with minimap-2.1 map-pb--N3 and resulting sam file was converted to bam and sorted (Li 2018; Li et al. 2009). Details of raw long reads used are provided in **Supplementary Table S4**. Three structural variant (SV) callers – sniffles-2.0, cuteSV-1.0.13, and svim-2.0 were used for SV detection. All three callers were run with default parameters but mapQ required to be >50 and minimum read support required adjusted for each strain by sequencing coverage i.e., 5 reads per 100X coverage(Jiang et al. 2020; Sedlazeck et al. 2018; Heller and Vingron 2019). Only insertions and deletions >30 bp and duplications and inversions of >10 kb were retained. Next, for each caller, biallelic SVs with precise mapping were selected and merged from 8 samples using Survivor (Jeffares et al. 2017) and only simple SVs (non-complex and unambiguous) were used. Summary of filtered and raw SV calls are in Supplementary Table 3.

Genotyping for merged SVs for each caller was performed by cuteSV-1.0.13 with min_support set as 3 reads. Genotyped calls that were supported by two SV callers or more, for which intra-strain allele frequency (AF) could be determined were then filtered and their SV length, AF, and read support averaged from results of SV callers using bedtools *merge* function. Additionally, overlapping SVs of same type with length difference by >20% or 500 bp were treated as independent events, otherwise collapsed as a same SV event. 71% of all simple SVs filtered were detected by at least two callers and 44.5% detected by all three callers. Next, *D. melanogaster* SVs were polarized by comparison to their absolute presence or absence in *D. simulans* and *D. sechellia*. Any SVs with conflicted calls between these two sister species were ignored. Filtered and polarized SV calls are reported in **Supplementary Table S5**.

### Structural variant parsing and enrichment analyses

Filtered SVs of insertions and deletions class were used for parsing for further analysis. The vast majority of SVs are expected to have extremely low frequency of <0.1, which reflects the general deleterious nature of SVs. Since raw long read data used from published studies were from pooled sequencing of ∼200 flies for DSPR strains and 60-80 flies for iso-1, only SVs with intra-strain AF of 0.2 or more were considered, to enrich for germline SVs. SV enrichment analysis was carried out using *poverlap* (https://github.com/brentp/poverlap) with 1000 bootstraps of random shuffling. Mean expected overlap counts against piC coordinates were compared to expected overlap.

### *de novo* TE annotation

To create a comprehensive and accurate representative TE library representing the TE insertions contained in the 8 strains, *de novo* TE annotation was conducted using several computational tools. Summary of all major steps is presented in a flow-chart in **Fig. 4A**. Briefly, canonical FlyBase TE consensus sequences were curated for each strain to include only TEs that best represent the TE insertion landscape of each strain using RepeatMasker-4.1.0 results(Smit 1999; Larkin et al. 2021). FlyBase TE families with at least 3 copies of >200 bp and <1% divergence was retained. This resulted in a reference TE library for each strain, averaging ∼110 TE families. Next, RepeatModeler2 was run on the 7 DSPR genomes (Chakraborty et al. 2019) and reference iso-1 strain(Hoskins et al. 2015) followed by removal of non-TE repeats like tRNA, satellites, rRNA etc., as well as TE sub-families using bash scripts (Flynn et al. 2020). Next, identification of novel TE families absent in the FlyBase TE library was carried out using the 80-80-80 rule (Wicker et al. 2007; Quesneville et al. 2005). TE consensus fragments from RepeatModeler2 library that passed the 80-80-80 rule using blastn (Camacho et al. 2009) alignments with reference TEs were removed and remaining likely “novel” ones were used to mask respective genome assemblies with RepeatMasker. From RepeatMasker results, all TE consensus fragments with at least 3 copies of >200 bp with <5% divergence from the consensus were retained as potential novel TE family fragments and combined into one compiled DSPR-library for all 8 strains.

### TE family and super-family enrichment analyses

RepeatMasker outputs of the comprehensive *D. melanogaster* TE library on respective genome assemblies of 8 strains was generated (Smit 1999). RM.out files were parsed, and insertions defragmented using scripts from https://github.com/4ureliek/Parsing-RepeatMasker-Outputs (Kapusta et al. 2017). Family-level and Superfamily-level enrichment analyses were conducted using the TE_analysis_Shuffle_bed.pl from https://github.com/4ureliek/TEanalysis (Kapusta et al. 2013). 1000 bootstraps were performed and only TE families with 10 or more total insertions were considered for enrichment analyses.

### Phylogenetic tree construction

Maximum likelihood (ML) trees were constructed for all defragmented iso-1 TE insertions for superfamilies and families reported in **Fig. 6** using the method described below. First, TE insertion sequences were extracted from the reference genome using bedtools *getfasta.* Sequences in the range of 200-500 bp were manually examined for further defragmentation if nearby insertions of the same family were present with non-overlapping sequences when compared to consensus. Sequences were then subjected to multiple sequence alignment using mafft v7.453 with the E-INS-i strategy and following parameters--adjustdirectionaccurately--maxiterate 1000. Next, ML trees were constructed using raxML-HPC with GTRGAMMA model (Stamatakis 2014). ML trees were then uploaded in R and terminal branch lengths calculated and visualized using the tidytree and ggtree R packages (Yu 2022; Yu et al. 2018).

### *de novo* TE insertion detection from long reads

Raw long reads utilized in SV detection was also used for *de novo* TE insertion analyses. Reads (>1 kb) were mapped to iso-1 reference genome (without Y-linked contigs and all contigs<20 kb) using minimap2.1 default parameters and resulting sam file converted to bam file, sorted and indexed using samtools (Li 2018; Li et al. 2009). TLDR, a *de novo* TE detection program, was run for each strain using comprehensive *D. melanogaster* TE library curated in this study (Ewing et al. 2020). Insertion calls with 2-3 different TE families were treated as nested or complex insertion, where each family was treated as an independent insertion. However, they were ignored if all insertions belong to the same superfamily classification to remove false-positive hits resulting from micro-homologies between related families. Nested calls were also ignored if they belonged to more than 3 families. TLDR was run with default parameters except for *--flanksize* 200 *--max_te_len* 15000 *-m* 2 and result insertions table filtered. TLDR calls retained for analyses by filtering for medianMapQ >20, TEmatch>80%, and intrasample frequency of >0.05 (supporting reads/empty reads). TE insertion enrichment analyses were performed using TE_analysis_Shuffle-bed.pl script (Kapusta et al. 2013). Filtered TLDR insertion calls for all 8 strains are reported in **Supplementary table S7**.

## Data availability

Small RNA libraries sequenced in this study are deposited in SRA under the project accession PRJNAXXXXX. Relevant source data for figures is listed in supplementary tables. Raw data files, TE consensus sequences and scripts are available at https://github.com/kerogens101/Dmel_piCs.

## Supporting information

Supplementary_TextAndFigCombined

## Acknowledgements

We thank Jullien Flynn for her assistance in RepeatModeler2 runs and consensus filtering and Michelle Stitzer for recommendations on structural variant analysis. We also thank Dan Barbash, Justin Blumenstiel and Andrew Grimson for helpful discussions on the analyses and members of the Clark and Feschotte laboratories for valuable feedback on the manuscript. This work was supported by grants from the National Institute of Health, R01-GM119125 to A.G.C and R35-GM122550 to C.F.

## Supplementary Tables

**Supplementary_Table_S1**. Summary statistics of sequencing depth and mapping of all small RNA libraries analyzed in this study.

**Supplementary_Table_S2**. All data downloaded and analyzed from public domain.

**Supplementary_Table_S3**. piRNA cluster (piC) annotations for each small RNA library per strain from *restrictive*, *proTRAC*, and master-List.

**Supplementary_Table_S4**. piC population frequency for each pipeline after remap to iso-1 genome (liftOver).

**Supplementary_Table_S5**. Collapsed, genotyped, filtered, and polarized structural variant calls. **Supplementary_Table_S6**. TE classification and activity determined from non-reference TE insertions. **Supplementary_Table_S7**. Filtered and collapsed TLDR non-reference TE insertion calls.

