## Supplementary_TextAndFigCombined for "Rapid evolution of piRNA clusters in the *Drosophila melanogaster* ovary"

### Supplementary methods

#### Small RNA library QC analysis

piRNA length profile of each library is plotted (**Supplementary Fig. S1A)** from FastQC-0.11.8 runs on fastq files after in-silico removal of reads mapping to rRNA, miRNA, tRNA, snoRNA and snRNAs. rRNA-derived reads in trimmed libraries was quantified by mapping reads to rRNA sequences extracted from iso-1 reference assembly using RefSeq annotations downloaded from FlyBase (**Supplementary Fig. S1B)**. Additionally, spearman correlation of trimmed reads mapped to TE consensus library are plotted for uniquely and multi-mapping reads from each library (**Supplementary Fig. S1 C,D**). Next, to determine the saturation in small RNA sequencing across libraries, reads were collapsed to retain only unique sequences by TBr2_collapse of NGS-toolbox (Rosenkranz et al. 2015) and each sequences' abundance quantified in non-collapsed library (Genzor et al. 2021). Abundance of unique putative (23-29nt) piRNA sequences are plotted by their cumulative contribution to the library depth for two replicates each of strains A1 and B6 (**Supplementary Fig. S1 E,F**).

#### piC annotation methods

The first method, we call "*restrictive",* is conducted by capturing piRNA-expressing domains based on uniquely mapping piRNAs across the genome in sliding 500-bp windows. To recover complete piCs using uniquely-mapping piRNAs only, low-mappability regions flanked by such domains were merged into a cluster and analyzed for 1U-bias and the density of unique piRNA sequences to remove false positives (Mohn et al. 2014). The *restrictive* piCs were then curated with stringent cutoffs in expression (>5 RPKM), density (8 hits/500 bp), and mappability of piRNA domains. While this method considers differential genomic mappability, it only uses uniquely mapped piRNAs which only comprise ~16-24% of total predicted piRNA reads in all libraries. Hence, piCs were also annotated using a second method, *proTRAC*, which utilizes all piRNA reads and normalizes piRNA expression by its mappability to uncover piCs even when there are virtually no uniquely mapping piRNAs (Rosenkranz and Zischler 2012). A third method, we call "*permissive*", is carried out similarly to *restrictive* but with lower cutoffs (2 RPKM and 2 hits/500 bp) to recover clusters from extremely low mappability regions and few uniquely mapping piRNAs.

#### Consensus TE library curation

Putative novel TE fragments in the compiled DSPR-library were then clustered using cd-hit c 0.8 -aL 0.05 -aS 0.8 -A 80 -sc 1 -T 0 -d 1000 -g 1 -M 0 to create a non-redundant library of putative novel TEs. Post-clustering, 176 novel TE fragments remained, 52 of which were then manually curated into TE consensus sequences that could be classified into TE subclass and superfamilies. However, all 176 fragments were utilized for RepeatMasker run presented in **Fig. 4B.** Manual curation of novel TE fragments consisted of three steps. First, as step 1, fragments were searched against the Dfam (Storer et al. 2021) database of *D. melanogaster*, with e-value increased to 0.1. If >50% of the sequence could not be matched to any Dfam entry, then in step 2, conserved domains were searched for such fragment consensus in the NCBI Conserved Domain v3.19 database with e-value increased to 0.1 (Lu et al. 2020). If a conserved domain or domains was found, for example – RNaseH or Integrase core, the consensus was retained in the library. In step 3, the fragment consensus name was changed to reflect the closest related family or superfamily identified for that consensus. Details of the comprehensive TE library classification is provided in **Supplementary Table S6** and library consensus provided in GitHub repository.

### Supplementary Fig. 1

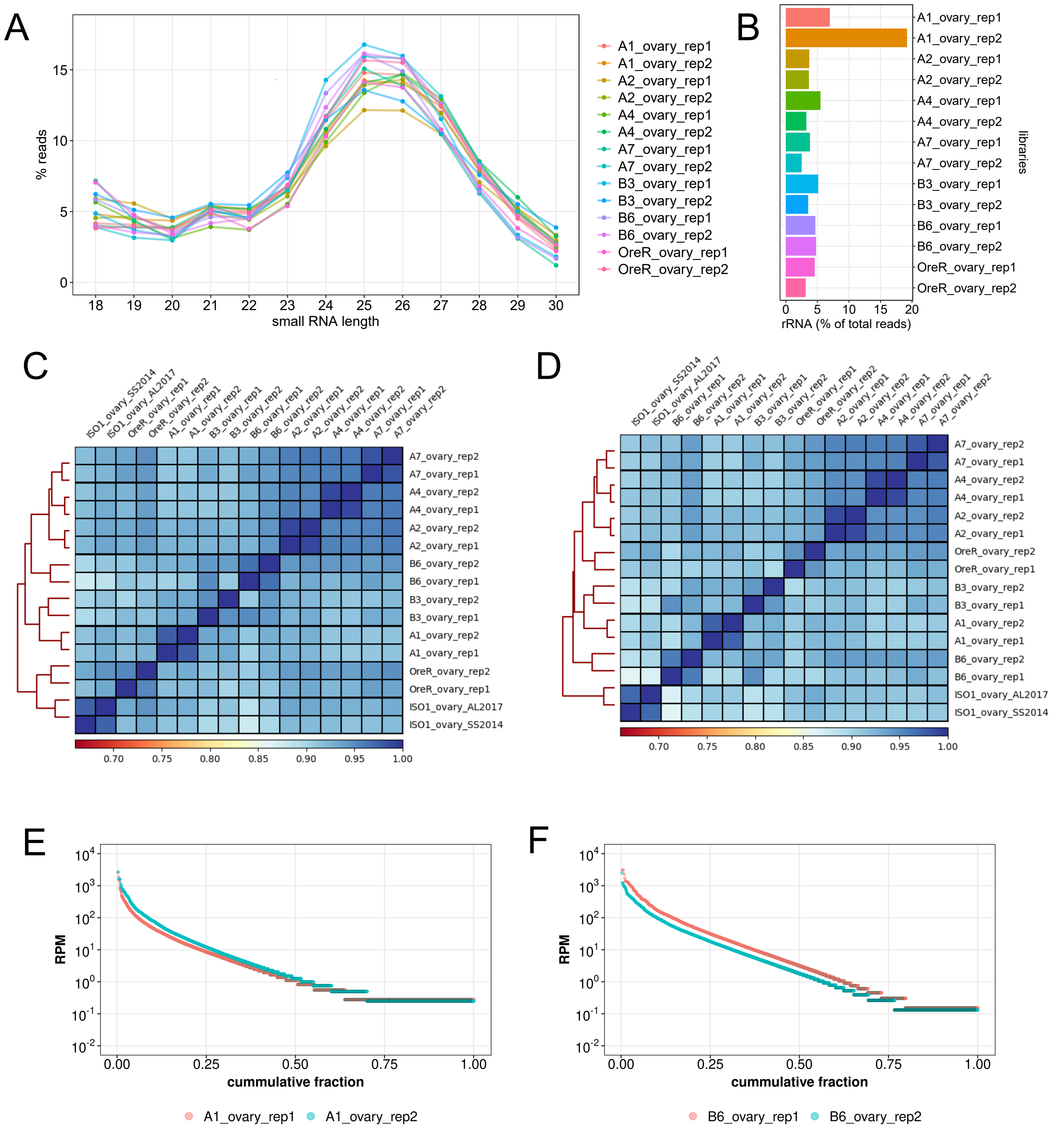


**Supplementary_Fig_S1. Small RNA library QC results.** *(A)* Small RNA read length profile after processing and removal of miRNA, tRNA, rRNA and snoRNAs. *(B)* Percentage of rRNA derived reads in small RNA libraries. *(C&D)* Spearman correlation plot of TE-derived small RNA expression in libraries using unique and multi mapping approaches respectively. *(E&F)* Comparison of cumulative sequence distribution of piRNAs in libraries to evaluate saturation of sequence space.

### Supplementary Fig. 2

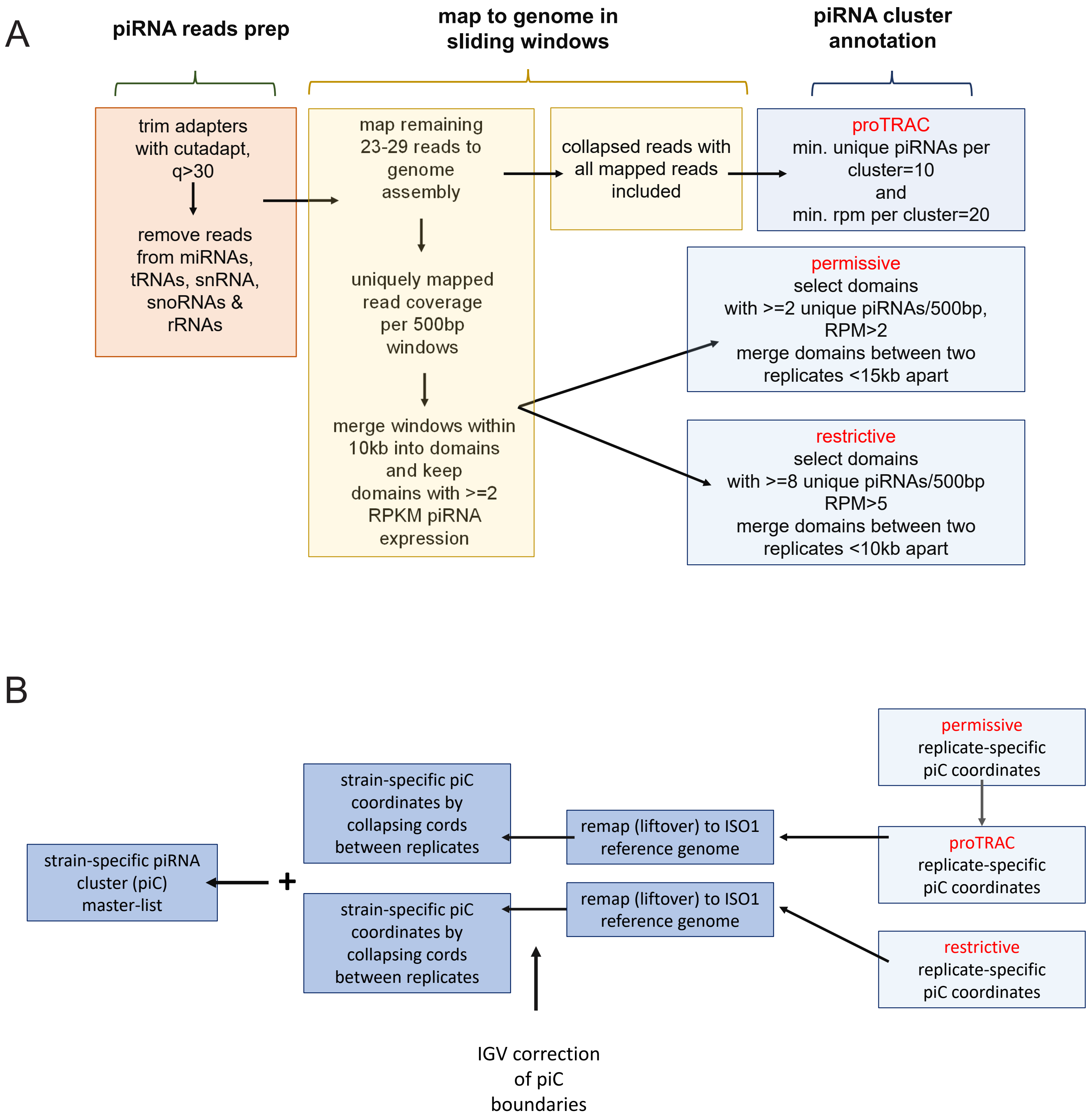


**Supplementary_Fig_S2. piRNA cluster (piC) annotation.** *(A)* Summary of the piC annotation pipeline using uniquely, and multi-mapping reads from each library independently. *(B)* Outline of steps in curation of master-list of piC annotations for each DSPR strain from restrictive and proTRAC method.

### Supplementary Fig. 3

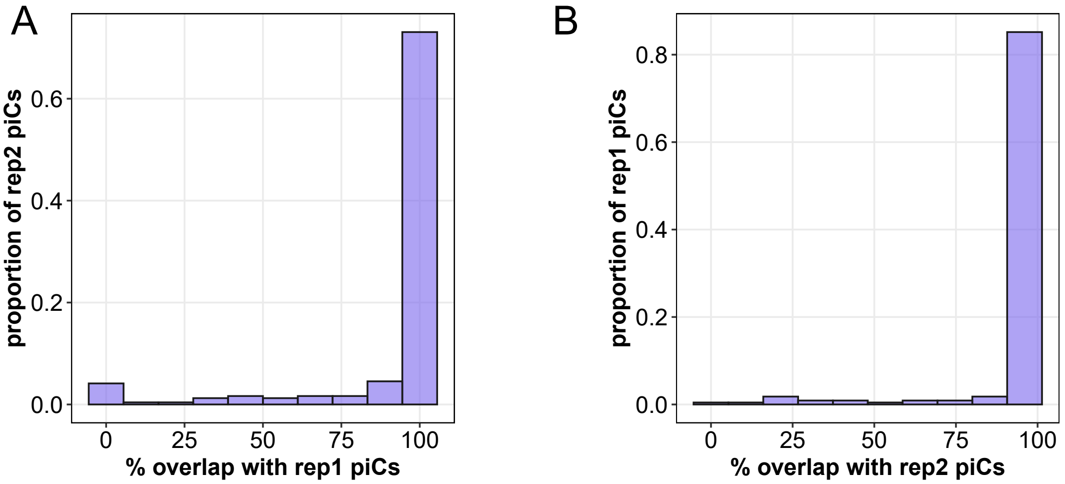


**Supplementary_Fig_S3. Reproducibility of piC annotations.** *(A&B)* Histogram showing proportion of piCs that overlap 0-100% of their respective length in annotations between two biological replicates of strain A1 small RNA libraries.

Srivastav et al, 2023

### Supplementary Fig. 4

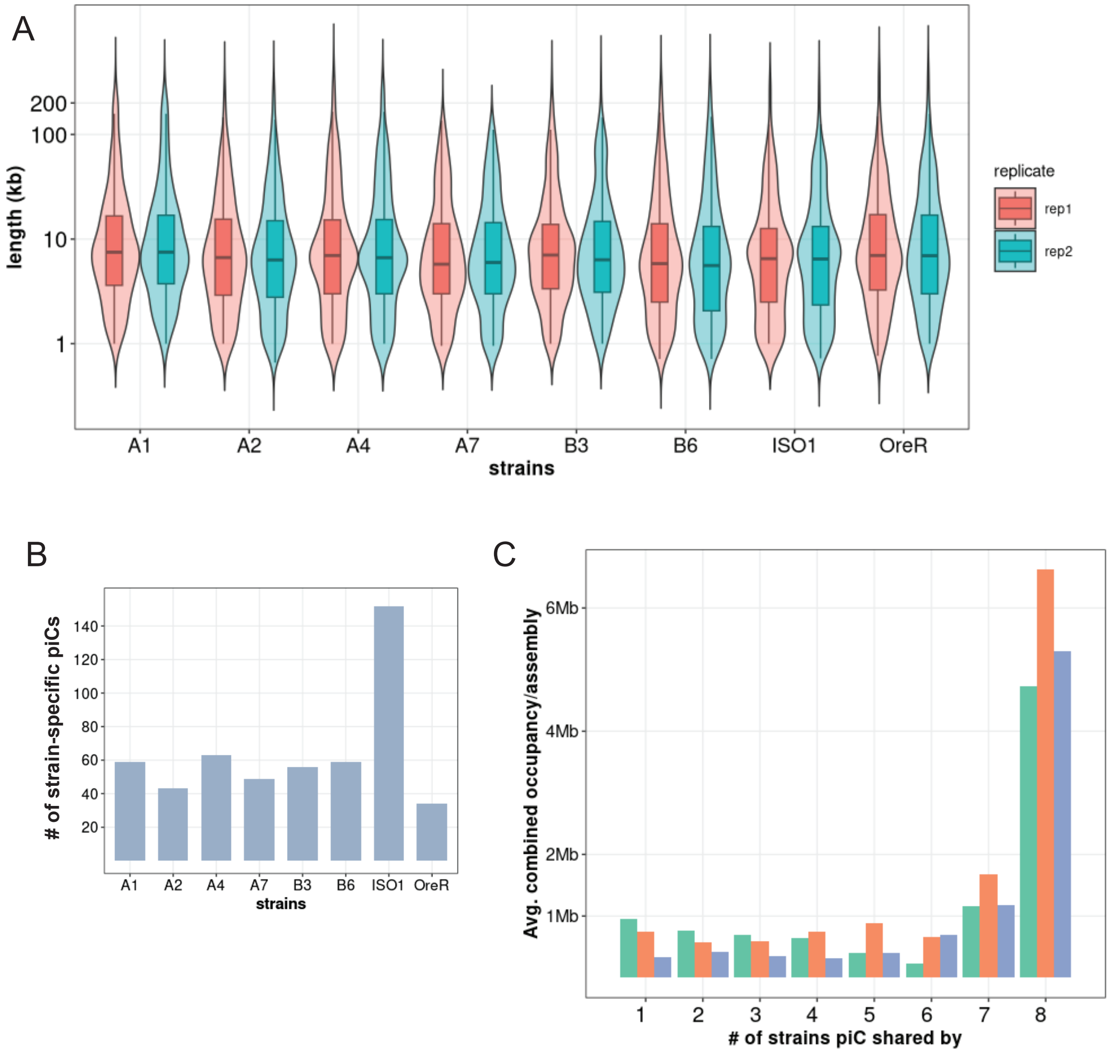


**Supplementary_Fig_S4.** ***piC length and cumulative size metrics of predicted piCs.*** *(A)* Length distribution of piCs predicted independently from each small RNA library. *(B)* Count of strain-specific piCs in the master-list from each strain. Due to liftOver/remapping, ~50% of piCs are lost from non-reference strains. *(C)* Cumulative piC size from each annotation pipeline and grouped by sharing of piCs among strains.

### Supplementary Fig. 5

A

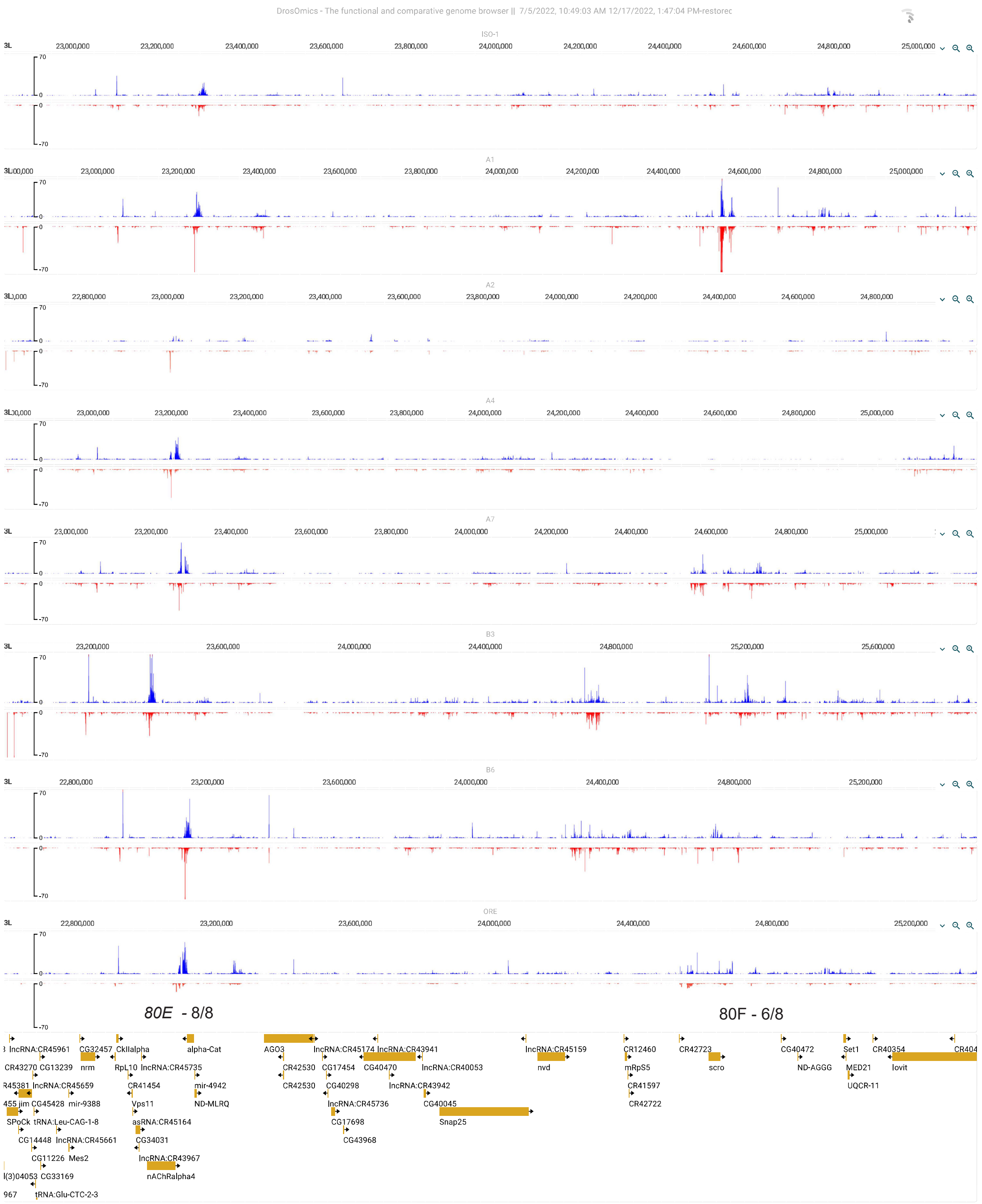


**Supplementary_Fig_S5. Additional examples of piCs with varying activity among strains.** *(A)* DrosOmics genome browser shot of two moderate to highly expressed piCs – 80E and 80F separated by multiple genes including *Ago3*. Numbers next to piC name indicates how many strains that piC is detected in by restrictive and proTRAC pipelines. piRNA coverage presented is average RPM from two replicate small RNA libraries per strain mapped to respective genome assemblies. *(B)* DrosOmics genome browser shot of two moderate expressed genic piCs – associated with Trypsin genes and *eEF1alpha1* separated by multiple genes including *Ago3.* Numbers next to piC name indicates how many strains that piC is detected in by restrictive and proTRAC pipelines. piRNA coverage presented is average RPM from two replicate small RNA libraries per strain mapped to respective genome assemblies.

Srivastav et al, 2023

### Supplementary Fig. 5

B

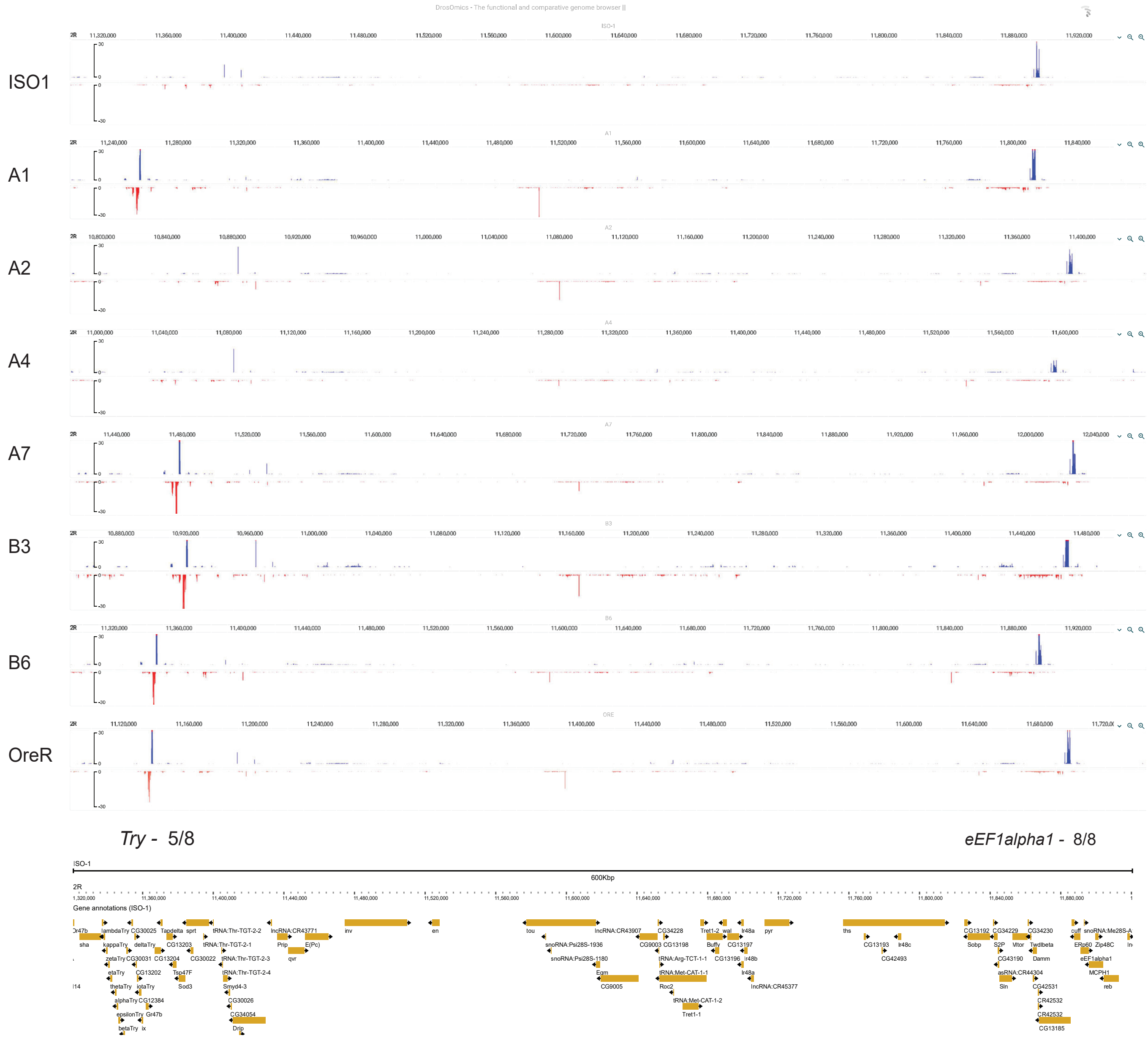

### Supplementary Fig. S6

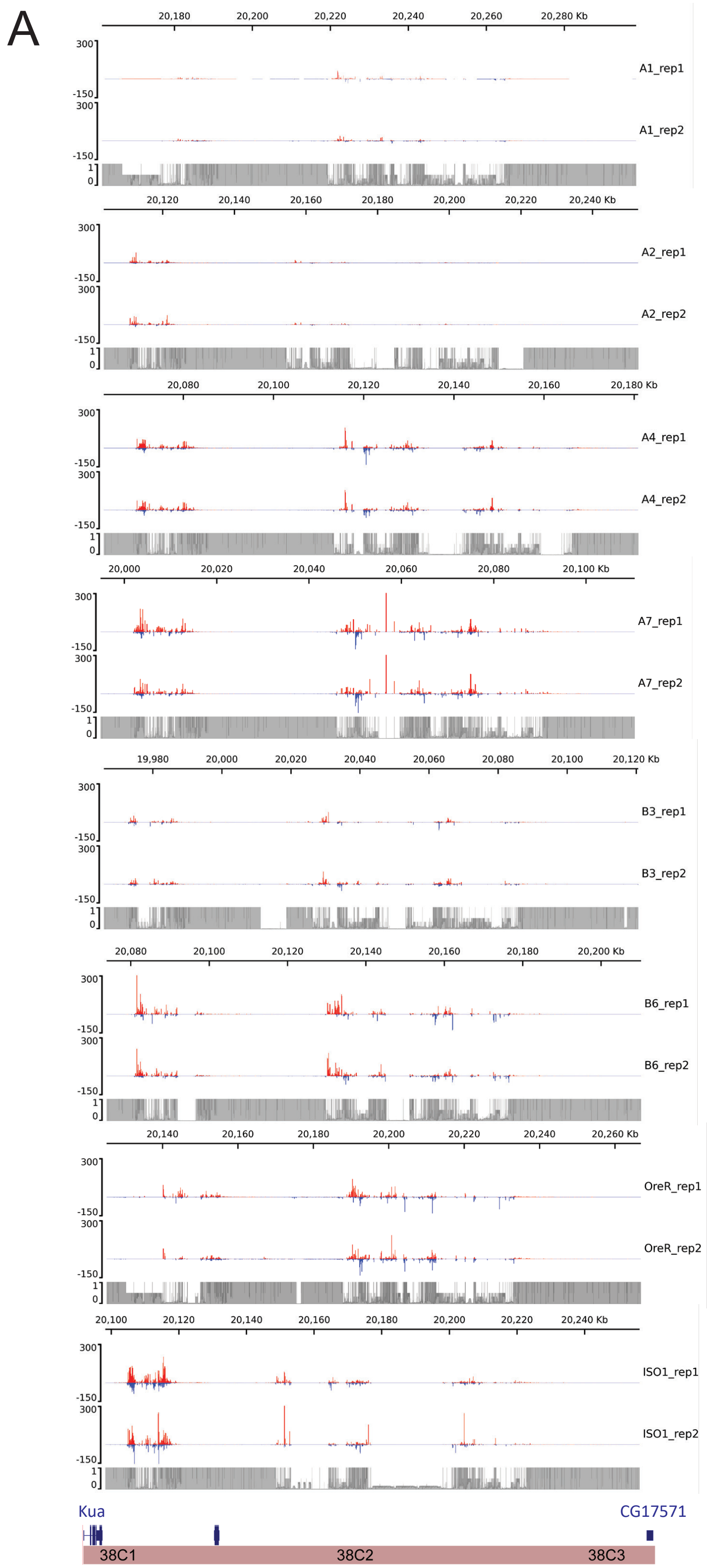


**Supplementary_Fig_S6. Additional example of natural variation in large highly expressed piC activity.** *(A)* Uniquely mapping piRNA expression profiles of *38C* piC for the 8 strains with two small RNA library replicates. Expression values are in reads per million (RPM) for 100 bp bins. Mappability scores (0-1) is shown for 100 bp bins of each respective *38C* genomic assembly.

### Supplementary Fig S7

Srivastav et al, 2023

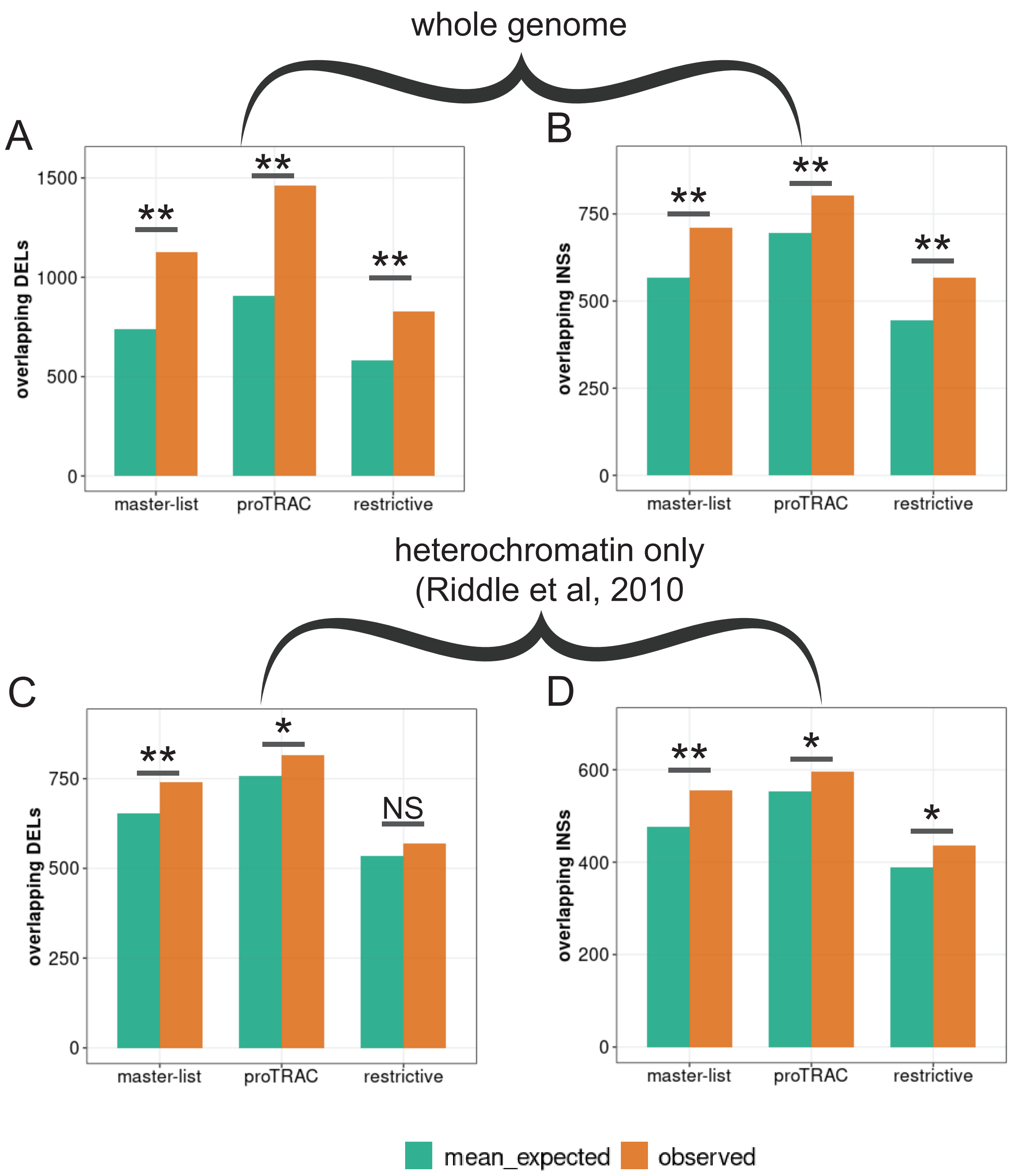


**Supplementary_Fig_S7. Comparisons of INDEL enrichment across whole genome and heterochromatin only to the piC regions from master-list.** *(A&B)* Bar plots displaying observed and mean expected values of INDELs overlapping piCs when piCs are shuffled across whole genome. *(C&D)* Bar plots displaying observed and mean expected values of INDELs overlapping piCs when piCs are shuffled only within heterochromatin boundaries.

Srivastav et al, 2023

### Supplementary_Table_S1

| sample | total reads (in millions) | piRNA reads (in millions) | rRNA reads (in millions) | %rRNA contamination | source |
|---|---|---|---|---|---|
| A1_ovary_rep1 | 25.73 | 13.7 | 0.47 | 1.83% | this study |
| A1_ovary_rep2 | 17.17 | 8.65 | 0.918 | 5.35% | this study |
| A2_ovary_rep1 | 23.66 | 11.33 | 0.233 | 0.98% | this study |
| A2_ovary_rep2 | 70.05 | 33.95 | 0.645 | 0.92% | this study |
| A4_ovary_rep1 | 20.67 | 10.94 | 0.333 | 1.61% | this study |
| A4_ovary_rep2 | 26.09 | 12.66 | 0.219 | 0.84% | this study |
| A7_ovary_rep1 | 18.77 | 9.3 | 0.196 | 1.04% | this study |
| A7_ovary_rep2 | 19.81 | 10.49 | 0.136 | 0.69% | this study |
| B3_ovary_rep1 | 9.11 | 5.29 | 0.122 | 1.34% | this study |
| B3_ovary_rep2 | 14.29 | 6.9 | 0.132 | 0.92% | this study |
| B6_ovary_rep1 | 20.41 | 11.42 | 0.253 | 1.24% | this study |
| B6_ovary_rep2 | 25.1 | 13.35 | 0.32 | 1.27% | this study |
| ISO1_ovary_SS2014 | 15.32 | 7.43 | 0.869 | 5.67% | Shipz S, et al 2014 |
| ISO1_ovary_AL2017 | 50.26 | 33.52 | 0.967 | 1.92% | Asif-Laidin A, et al 2017 |
| OreR_ovary_rep1 | 44.52 | 21.1 | 0.563 | 1.26% | this study |
| OreR_ovary_rep2 | 22.2 | 11.65 | 0.187 | 0.84% | this study |
